# Network asynchrony underlying increased broadband gamma power

**DOI:** 10.1101/2020.08.26.265439

**Authors:** Nicolas Guyon, Leonardo Rakauskas Zacharias, Eliezyer Fermino de Oliveira, Hoseok Kim, João Pereira Leite, Cleiton Lopes-Aguiar, Marie Carlén

**Affiliations:** Department of Neuroscience, Karolinska Institutet, 17177 Stockholm, Sweden; Department of Neuroscience and Behavioral Sciences, Ribeirão Preto Medical School, Universidade de São Paulo, 14049-900 Ribeirão Preto, Brazil; Center for Mathematics, Computing and Cognition - Universidade Federal do ABC - 09606-070 São Bernardo do Campo, Brazil; Dominick P Purpura Department of Neuroscience, Albert Einstein College of Medicine, Bronx, NY 10461, USA; Núcleo de Neurociências, Department of Physiology and Biophysics, Institute of Biological Sciences, Universidade Federal de Minas Gerais, 31270-901 Belo Horizonte, Brazil; Department of Biosciences and Nutrition, Karolinska Institutet, 14183 Huddinge, Sweden

## Abstract

Synchronous activity of cortical inhibitory interneurons expressing parvalbumin (PV) underlies the expression of cortical gamma rhythms. Paradoxically, deficient PV inhibition is associated with increased broadband gamma power. Increased baseline broadband gamma is also a prominent characteristic in schizophrenia, and a hallmark of network alterations induced by N-methyl-D-aspartate receptor (NMDAR) antagonists such as ketamine. It has been questioned if enhanced broadband gamma power is a true rhythm, and if rhythmic PV inhibition is involved or not. It has been suggested that asynchronous and increased firing activities underlie broadband power increases spanning the gamma band. Using mice lacking NMDAR activity specifically in PV neurons to model deficient PV inhibition, we here show that local LFP (local field potential) oscillations and neuronal activity with decreased synchronicity generate increases in prefrontal broadband gamma power. Specifically, reduced spike time precision of both local PV interneurons and wide-spiking (WS) excitatory neurons contribute to increased firing rates, and spectral leakage of spiking activity (spike “contamination”) affecting the broadband gamma band. Desynchronization was evident at multiple time scales, with reduced spike-LFP entrainment, reduced cross-frequency coupling, and fragmentation of brain states. While local application of S(+)-ketamine in wildtype mice triggered network desynchronization and increases in broadband gamma power, our investigations suggest that disparate mechanisms underlie increased power of broadband gamma caused by genetic alteration of PV interneurons, and ketamine-induced power increases in broadband gamma. Our studies, thus, confirm that broadband gamma increases can arise from asynchronous activities, and demonstrate that long-term deficiency of PV inhibition can be a contributor.

## Introduction

Cortical fast-spiking (FS) inhibitory interneurons expressing PV (FS-PV) regulate excitatory spike generation and timing, and help maintain an appropriate balance between inhibition and excitation in cortical circuits (Hu et al., 2014; Sohal and Rubenstein, 2019). Particularly, perisomatic PV inhibition is central to the coordinated interaction of excitation and inhibition that underlies the emergence of gamma oscillations, synchronized rhythmic high-frequency (30-80 Hz) patterns of electrical activity observed in the LFP (Buzsáki and Wang, 2012). Cortical PV interneurons receive NMDAR-dependent excitatory input from pyramidal neurons, and mutant mice lacking NMDAR activity specifically in PV-expressing neurons (PV-Cre/NR1f/f mice) display increased power of spontaneous gamma oscillations, i.e., increased power across the 30-80 Hz frequency band (Korotkova et al., 2010; Carlén et al., 2012; Gandal et al., 2012; Billingslea et al., 2014). LFP oscillations spanning across the 30-80 Hz band will hereafter be referred to as *broadband gamma oscillations*. Increased spontaneous broadband gamma oscillations have also been demonstrated in other mouse models of PV neuron dysfunctions (del Pino et al., 2013; Cho et al., 2015), and in individuals with schizophrenia (Mathalon and Sohal, 2015; Grent-’t-Jong et al., 2018). Further, administration of NMDAR antagonists such as ketamine (Pinault, 2008; Hakami et al., 2009; Kulikova et al., 2012; Caixeta et al., 2013; Picard et al., 2019; Lopes-Aguiar et al., 2020; Mahdavi et al., 2020; McNally et al., 2020), or MK-801 (Carlén et al., 2012; Molina et al., 2014; Hudson et al., 2020), is associated with increased power in spontaneous broadband gamma oscillations in both anesthetized and awake rodents, and also in humans (Rivolta et al., 2015). Importantly, lack of NMDAR in PV neurons blunts this response (Carlén et al., 2012; Picard et al., 2019; Hudson et al., 2020), confirming that cortical PV interneurons are a central target of NMDAR antagonists. However, it is unclear if pharmacologically-induced increases in broadband gamma oscillations are distinct from the increased spontaneous broadband gamma oscillations observed in genetic models of deficient PV inhibition, e.g. mice with lack of NMDAR activity in PV neurons (PV-Cre/NR1f/f mice).

Computational modeling has suggested that decreased excitability of PV interneurons due to an absence of slow excitatory NMDA currents would lead to reduced sensitivity to excitatory inputs, and the requirement of a more synchronous excitatory input for PV firing (Carlén et al., 2012). This “suppression boundary” mechanism (Börgers and Kopell, 2005) is modelled to enhance gamma oscillations over a broad frequency range (Carlén et al., 2012; Jadi et al., 2016). Independently, optogenetic experiments have demonstrated that increased excitability of excitatory neurons in the medial PFC (mPFC) increases spontaneous broadband gamma power (Yizhar et al., 2011), which has led to the suggestion that broadband increases in spontaneous gamma power can reflect an overall increased activity in cortical circuits (Sohal and Rubenstein, 2019). In line with this suggestion, systemic administration of the NMDAR antagonist MK-801 has demonstrated an association between increased broadband gamma (here 30-90 Hz) power and increased firing in the mPFC of rats (Molina et al., 2014). Importantly, the spike trains were found to be disorganized, and the synchronization of action potential firing decreased (Molina et al., 2014). This supports the view that increased gamma power might not (always) be a rhythm (a genuine oscillation) but rather a result of increases in uncorrelated synaptic potentials and neuronal discharges (Uhlhaas and Singer, 2015, and see references in Buzsáki and Wang, 2012), i.e., spontaneous broadband gamma oscillations could be fundamentally different from PV interneuron-generated gamma synchrony (Sohal and Rubenstein, 2019). Of relevance, both modelling and experimental findings show that dysfunction of PV inhibition results in increased variability of cortical excitatory activity (Carlén et al., 2012; Jadi et al., 2016). The mechanism(s) by which asynchronous neuronal activity caused by deficient PV inhibition could manifest as increased broadband gamma oscillatory activity has not been established.

Cortical gamma oscillations can be persistent or nested to slower (< 4 Hz) oscillations, depending on the ongoing cortical state. For instance, during wakefulness and *rapid-eye movement* (REM) sleep, i.e. cortical states marked by lower LFP amplitude and faster frequencies (> 30 Hz), gamma oscillations are persistent over time. During non-REM (NREM) sleep, a cortical state characterized by cyclic alternations between silent (DOWN) and increased (UP) neuronal activity, gamma oscillations are nested to the delta oscillations (0.5-4 Hz) (Steriade, 2006; Harris and Thiele, 2011; McKenna et al., 2017). This pattern of gamma modulation is also observed under urethane anesthesia, which is characterized by cyclic alternations between cortical states that resemble NREM sleep oscillations (deactivated states) and REM sleep oscillations (activated states) (Clement et al., 2008). Systemic administration of ketamine alters the cyclic alternations between deactivated and activated states, and the coupling between delta and gamma oscillations in the mPFC in urethane anesthetized rats (Lopes-Aguiar et al., 2020). Importantly, PV inhibition is suggested to regulate state transitions (Zucca et al., 2017). How deficiency in PV inhibition affects cortical states is at large unknown.

To directly investigate a connection between deficient PV inhibition, increased power of broadband gamma oscillation, and asynchronous neuronal activity, we recorded cortical LFP and single-unit activity in mice lacking NMDAR activity in PV neurons (PV-Cre/NR1f/f mice). The recordings were targeted to the PFC, as alterations of PV interneurons have been linked to prefrontal dysfunction and cognitive deficits in psychiatric disorders such as schizophrenia (Lewis et al., 2012). Further, to enable examination of how lack of NMDAR activity specifically in PV neurons affects cortical states, and the coupling of gamma oscillations to slower oscillations, the recordings were conducted during urethane anesthesia. Our experiments revealed altered mPFC state transitions, with fragmentation of the deactivated states, and increased temporal variability in the firing of both excitatory neurons and PV interneurons in mice with lack of NMDAR activity in PV neurons. Additionally, PV-Cre/NR1f/f mice displayed DOWN and UP state events with increased power in the broadband gamma and high-frequency oscillations (HFOs; 100-150 Hz) bands, marked by increased mPFC firing rates, and our analyses show that this increased high-frequency power in part represents spectral leakage of spiking activity.

For comparative investigations of the effects of local NMDAR antagonists on prefrontal oscillatory and neuronal activity, and their dependence on NMDAR in PV interneurons, S(+)-ketamine was locally applied into the mPFC during the course of the recordings. Application of ketamine triggered a state with desynchronized LFP oscillations and increased broadband gamma power in (control) mice with intact NMDAR activity in PV neurons. In contrast, ketamine did at large not alter network activities in PV-Cre/NR1f/f mice, supporting the view that ketamine acts on NMDAR in inhibitory PV interneurons. Moreover, ketamine induced increased firing of prefrontal neurons and shifted the spike entertainment towards higher LFP frequencies in mice with intact NMDAR activity in PV neurons, effects absent in mice lacking NMDAR activity in PV neurons. Together, our investigations demonstrate that long-term deficiency of PV inhibition (here modelled by absence of NMDAR activity specifically in PV neurons) can result in increased and asynchronous neuronal activity, contributing to increased broadband gamma power of LFP oscillations by spectral leakage, and altered spike-LFP entrainment and state transitions. Our analyses, further, suggest that disparate mechanisms underlie increased power of broadband gamma caused by genetic alteration of PV interneurons, and ketamine-induced power increases in broadband gamma, respectively.

## Results

To establish how dysfunction of PV inhibition affects synchrony in cortical networks, mice lacking NMDAR activity in PV neurons were used. Deletion of the NMDAR subunit NR1 ablates all NMDAR activity in the affected neuron (Forrest et al., 1994), and studies in transgenic mice has shown that NR1 deletion in PV neurons leads to cognitive dysfunction and aberrations in cortical network activity with direct relevance to the symptomatology in disorders such as schizophrenia (Carlén et al., 2012; Gandal et al., 2012; Hudson et al., 2020; see review by Ju and Cui, 2015). Mice expressing Cre recombinase in PV neurons (PV-Cre mice) were crossed with mice carrying floxed alleles of the NR1 subunit (NR1f/f mice), generating PV-Cre/NR1f/f mice. PV-Cre mice were used as controls. To allow for light-activation and identification of the activity of mPFC PV interneurons in the electrophysiological recordings (opto-tagging; Kim et al., 2016, 2020), an adeno-associated virus (AAV) vector with Cre-dependent expression of channelrhodopsin-2 (ChR2) fused to mCherry (AAV-DIO-ChR2-mCherry) was unilaterally targeted to the mPFC (Cardin et al., 2009) (PV-Cre/NR1f/f, n = 3, PV-Cre, n = 4; **Figs. S1A-C**). To control for any optogenetics artifacts, four PV-Cre mice were injected with an AAV with Cre-dependent expression of the fluorophore eYFP (AAV-DIO-eYFP; **Figs. S1A-C**). Electrophysiological LFP recordings were conducted under urethane anesthesia five to seven weeks after viral injections using a silicon probe (four shanks with two tetrodes each) targeted to the prelimbic area (PL) in the mPFC (**Figs. S1D, E**). Urethane anesthesia is a validated model for testing of mechanistic hypotheses regarding sleep-like brain oscillations (Hauer et al., 2019), and has been particularly employed for investigation of the effects of NMDAR antagonists on oscillatory patterns in different brain circuits (Kiss et al., 2013; Lopes-Aguiar et al., 2020). Use of an anesthetized preparation circumvents the increases in gamma power associated with hyperlocomotion elicited by administration of NMDAR antagonists in awake mice (Hakami et al., 2009).

### Mice with lack of NMDAR activity in PV neurons display altered cortical states and oscillatory patterns

Urethane anesthesia displays spontaneous fluctuations between a deactivated state, characterized by slow oscillations (0.5-2 Hz) that resembles NREM sleep, and an activated state, characterized by faster oscillations (> 3 Hz), mirroring aspects of rapid eye movement (REM) sleep (Pagliardini et al., 2013). For characterization of the state dynamics over time, we divided the continuous LFP recordings (∼50 min) into 30 seconds epochs, and integrated the power for the 0.5-2 Hz and 3-45 Hz frequency bands for each epoch. Deactivated and activated brain states were classified using a Gaussian Mixture Model (GMM) clustering approach using the power of the 0.5-2 Hz and 3-45 Hz bands (**Figs. 1A-D**). The mPFC activity in PV-Cre mice displayed typical rhythmic transitions between activated and deactivated states (**Fig. 1A**), with the activated state being characterized by high power in the 3-45 Hz frequency band and the deactivated state by high-power in the slow oscillation bandwidth (0.5-2 Hz) (**Fig. 1D**). The strong presence of slow oscillations specifically during deactivated states can be observed in the spectrogram using short-time Fourier transform (**Fig. 1A**). In between the two state types, a transition period was identified which exhibited intermediate electrographic characteristics (**Fig. 1C**). These short transition periods were not further analyzed. While deactivated and activated states were present, and alternated, in PV-Cre/NR1f/f mice (**Figs. 1B, C**), the lack of NMDAR activity in PV neurons affected the state dynamics. The proportion of time spent in deactivated vs activated states, respectively, did not differ between PV-Cre/NR1f/f and PV-Cre mice, but, the deactivated states were significantly more frequent in PV-Cre/NR1f/f mice than in PV-Cre mice (**Figs. 1E, F**) and, thus, had a significant shorter mean duration than the deactivated states in PV-Cre mice (**Figs. 1G-I**). No apparent difference in either frequency or duration of the activated states was found between PV-Cre/NR1f/f and PV-Cre mice (**Figs. 1E-G**). In agreement with our results, sleep fragmentation has mainly been observed during NREM stages in both schizophrenia patients and animal models (Chouinard et al., 2004; Phillips et al., 2012).

**Figure 1.**
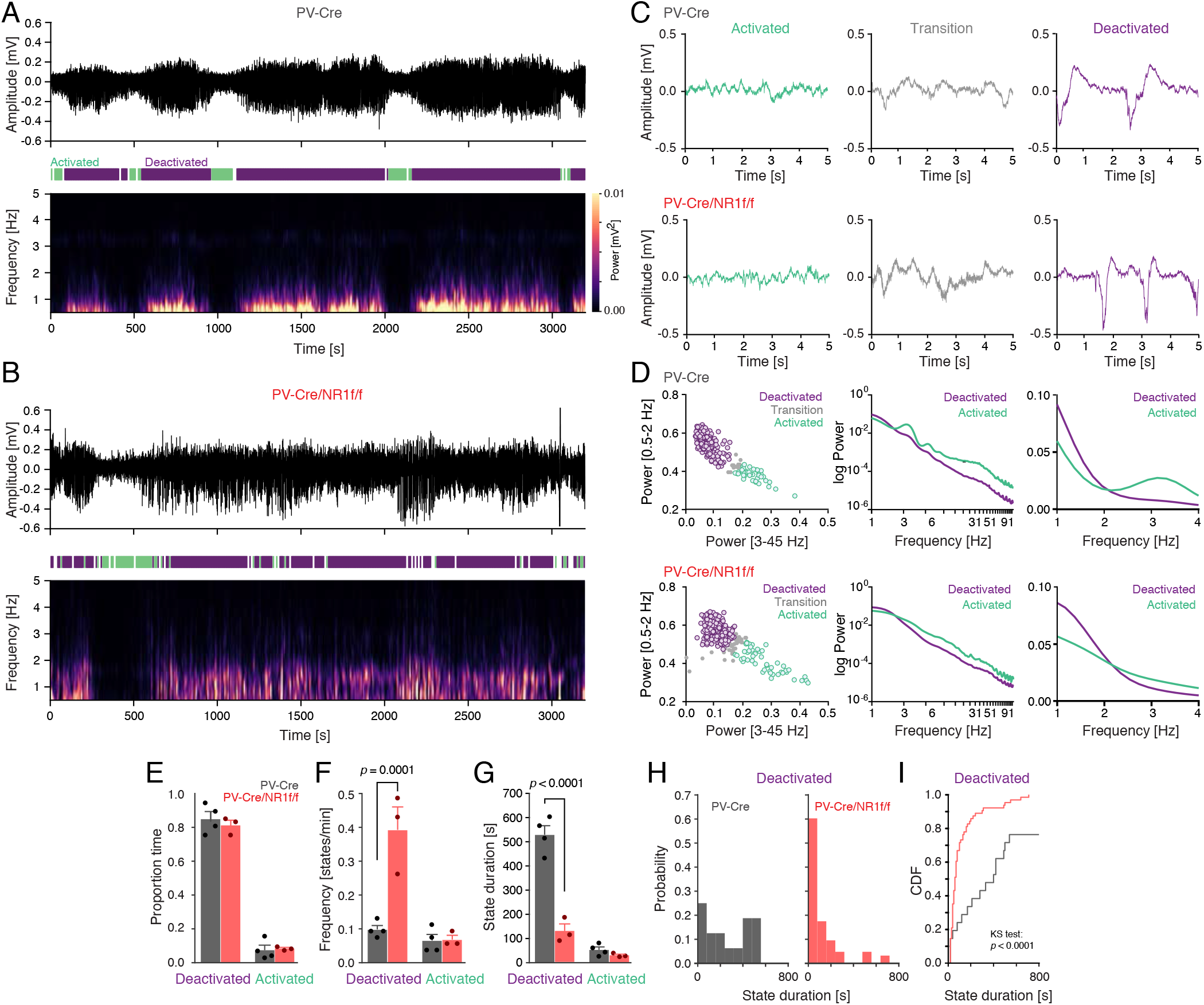
Mice with lack of NMDAR activity in PV neurons display altered cortical states. **A-B**) Representative mPFC LFP oscillations (3200 s) recorded under urethane anesthesia in a PV-Cre (**A**), and a PV-Cre/NR1f/f (**B**) mouse, respectively. Top: Raw LFP; middle: color-coded outline of detected activated (green), deactivated (purple) and transition states (white); bottom: Spectrogram (0-5 Hz) of the LFP trace in the top panel. **A)** Mice with intact NMDAR in PV neurons (PV-Cre) display typical cyclic transitions between deactivated (high amplitude low frequency (0.5-2 Hz) oscillations), and activated states (low amplitude higher frequency oscillations). **B)** PV-Cre/NR1f/f mice show less defined slow oscillations along with altered transition patterns between deactivated and activated states. **C)** Representative LFP traces (5 s) of activated (green), transition (grey), and deactivated states (purple), respectively, from a PV-Cre mouse (top) and a PV-Cre/NR1f/f mouse (bottom). **D)** Classification of brain states in PV-Cre mice (n = 4, top) and PV-Cre/NR1f/f mice (n = 3, bottom). Left: For detection of deactivated, activated, and transition (unclassified) states, a Gaussian mixture model was used for clustering of 30 s epochs (colored circles) of the LFP trace based on the power in the 0.5-2 Hz and 3-45 Hz frequency bands. Middle: Relative Power Spectrum Density (PSD, 1-91 Hz) of the LFP for the activated (green) and deactivated (purple) states. Right: Magnification of the relative PSD of the LFP in the 1-4 Hz frequency band. **E-I**) Comparison for the proportion (of total time), frequency (states/min), and duration, of deactivated and activated states in PV-Cre mice (n = 4) and PV-Cre/NR1f/f mice (n = 3). **E)** There is no difference in the proportion time spent in deactivated and in activated states between PV-Cre/NR1f/f and PV-Cre mice (F_(1,10)_ = 0.1908, *p* = 0.6715). Deactivated: PV-Cre: 0.85 ± 0.04; PV-Cre/NR1f/f: 0.81 ± 0.03; *p* = 0.8768; activated: PV-Cre: 0.07 ± 0.03; PV-Cre/NR1f/f: 0.08 ± 0.01; *p* > 0.9999. **F)** Mice lacking NMDAR in PV neurons (PV-Cre/NR1f/f) transition into deactivated states significantly more often than mice with intact NMDAR in PV neurons (PV-Cre mice) (F_(1,10)_ = 0.2252, *p* = 0.0008). Frequency (states/min) of deactivated: PV-Cre: 0.098 ± 0.012 states/min; PV-Cre/NR1f/f: 0.394 ± 0.068 states/min; *p* = 0.0001; activated: PV-Cre: 0.066 ± 0.018 states/min; PV-Cre/NR1f/f: 0.069 ± 0.013 states/min; *p* > 0.9999. **G)** The duration of the deactivated states is significantly shorter in mice lacking NMDAR in PV neurons (PV-Cre/NR1f/f) than in mice with intact NMDAR in PV neurons (PV-Cre) (F_(1,10)_ = 64.89, *p* < 0.0001). Deactivated: PV-Cre: 530.46 ± 37.31 s; PV-Cre/NR1f/f: 130.63 ± 29.09 s; *p* < 0.0001; activated: PV-Cre: 51.88 ± 11.79 s; PV-Cre/NR1f/f: 29.33 ± 4.44 s; *p* > 0.9999. **H)** Probability histogram of the duration of individual deactivated states in PV-Cre (gray) and PV-Cre/NR1f/f (red) mice. **I)** Lack of NMDAR in PV neurons results in significantly shorter deactivated states, as confirmed by comparison of the empirical cumulative distribution function (CDF) of the deactivated state durations between PV-Cre/NR1f/f and PV-Cre mice (*p* < 0.0001). Data are shown as mean ± SEM. For (**E-G**) Two-way ANOVA was used to assess significance, followed by a Bonferroni’s multiple comparisons test in case the ANOVA comparisons reached significance. For the cumulative distribution function in (**I**), the Kolmogorov-Smirnov test was used to assess significance.

Consistent with the literature (Carlén et al., 2012; Gandal et al., 2012; Jadi et al., 2016), lack of NMDAR activity in PV neurons resulted in increased LFP power over a broad frequency spectrum (**Fig. 2A**). This increase was specifically observed during the deactivated state, and particularly pronounced for the mPFC HFO frequency band, which showed a significant difference between PV-Cre and PV-Cre/NR1f/f mice (**Fig. 2B**). No significant power differences between PV-Cre and PV-Cre/NR1f/f mice were detected in any frequency band (0.5-150 Hz) during activated states (data not shown). Faster oscillations (> 30 Hz) can be modulated by different phases of low frequency LFP oscillations, an interaction suggested to regulate the spike probability and integrate activity across different temporal scales (Canolty and Knight, 2010). We therefore next investigated the modulation of faster oscillations by the phase of slower oscillations during the deactivated states. For this, we built phase-amplitude comodulation maps (comodulograms) (Tort et al., 2010) which identified that the strongest comodulation was found between low-delta frequency (0.5-2 Hz) phase, and broadband gamma and HFO (100-150 Hz) amplitude (**Figs. 2C, D**). Comparison of the Modulation Index (MI) revealed that the amplitude of both the high-frequency bands were significantly less modulated by the slow oscillation in the PV-Cre/NR1f/f mice than in PV-Cre mice (**Figs. 2E-H**). Taken together, the results suggest that intact NMDAR activity in PV neurons is required for proper temporal coordination of cortical network activity at multiple timescales, including coordination of cortical states (activated-deactivated states; minutes), and coordination between faster and slower oscillations of the LFP (seconds).

**Figure 2.**
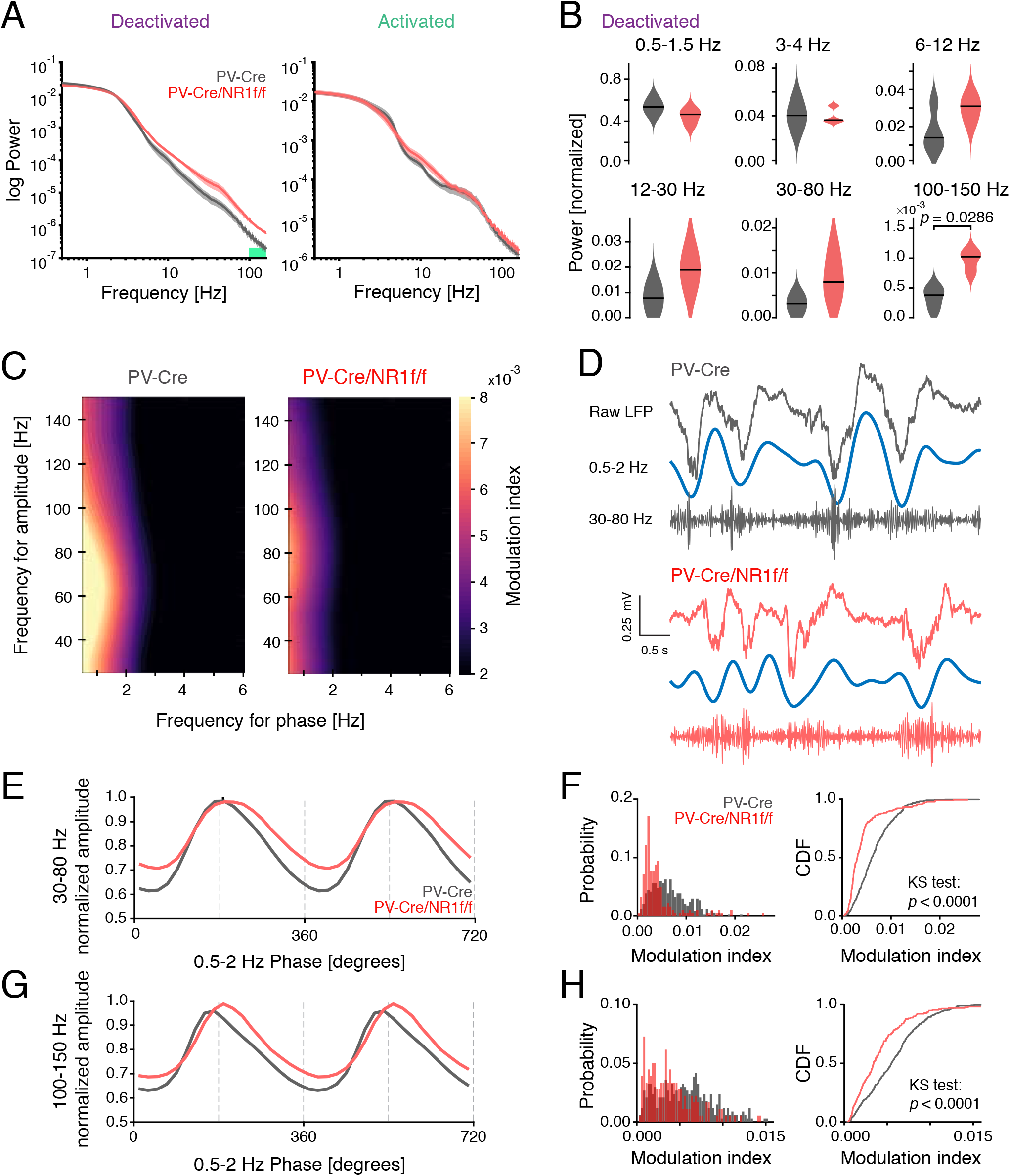
Mice with lack of NMDAR activity in PV neurons display increased broadband gamma power and decreased delta-phase modulation of broadband gamma and HFO amplitude. **A)** Mean PSD of the LFP for the PV-Cre (n = 4; gray) and PV-Cre/NR1f/f (n = 4; red) mice of deactivated (left) and activated (right) states. Green bar: frequency band (100-150 Hz) with significant power difference between PV-Cre and PV-Cre/NR1f/f mice. **B)** Comparisons of the integrated power at different frequency bands during deactivated states between PV-Cre and PV-Cre/NR1f/f mice. Low-delta (0.5-1.5 Hz): PV-Cre: 0.531; PV-Cre/NR1f/f: 0.462; *p* = 0.1143; high-delta (3-4 Hz): PV-Cre: 0.041; PV-Cre/NR1f/f: 0.037; *p* = 0.4286; theta (6-12 Hz): PV-Cre: 0.014; PV-Cre/NR1f/f: 0.031; *p* = 0.1143; beta (12-30 Hz): PV-Cre: 0.008; PV-Cre/NR1f/f: 0.0189; *p* = 0.0571; broadband gamma (30-80 Hz): PV-Cre: 0.003; PV-Cre/NR1f/f: 0.008; *p* = 0.1143; HFO (100-150 Hz): PV-Cre: 0.00038; PV-Cre/NR1f/f: 0.00102; *p* = 0.0286. **C)** Representative phase-amplitude comodulograms of a PV-Cre (left), and a PV-Cre/NR1f/f mouse (right), respectively, identifying that that the strongest comodulation was found between low-delta frequency (0.5-2 Hz) phase, and broadband gamma and HFO (100-150 Hz) amplitude. **D)** Representative unfiltered LFP trace (top) and filtered delta (0.5-2 Hz; middle) and broadband gamma (30-80 Hz; bottom) bands during a deactivated state in a PV-Cre (gray) and a PV-Cre/NR1f/f (red) mouse, respectively. **E)** Mean broadband gamma amplitude (normalized) at different phases of the low-delta cycles for PV-Cre (n = 4; gray) and PV-Cre/NR1f/f mice (n = 3; red), respectively. **F)** Left: Distribution of the modulation index probability between broadband gamma amplitude and low-delta phase in PV-Cre (n = 4; gray) and PV-Cre/NR1f/f mice (n = 3; red). Right: Comparison of the empirical cumulative distribution function of the modulation index shows that PV-Cre/NR1f/f mice have significantly more 30 s epochs of lower modulation index than PV-Cre (*p* < 0.0001), indicative of lower comodulation between broadband gamma amplitude and low-delta phase. **G)** Mean HFO amplitude (normalized) at different phases of the low-delta cycles for PV-Cre (n = 4; gray) and PV-Cre/NR1f/f mice (n = 3; red), respectively. **H)** Left: Distribution of the modulation index probability between HFO and low-delta phase in PV-Cre (n = 4; gray) and PV-Cre/NR1f/f mice (n = 3; red). Right: Comparison of the empirical cumulative distribution function of the modulation index shows that PV-Cre/NR1f/f mice have significantly more 30 s epochs of lower modulation index than PV-Cre mice (*p* < 0.0001), indicative of lower comodulation between HFO amplitude and low-delta phase. For (**A**), lines: mean; shaded area: SEM. For (**B**), black lines: median. For (**B**), two-tailed unpaired t-test was used to assess significance if data passed the D’Agostino & Pearson normality test, if not, the Mann Whitney test test was used. For the cumulative distribution function (**F, H**), the Kolmogorov-Smirnov test was used to assess significance.

### Mice with lack of NMDAR activity in PV neurons display altered DOWN and UP states

Under urethane, the deactivated brain state is dominated by slow oscillations, constituted by cyclic alternations between DOWN and UP states. During DOWN states, a marked period of populational hyperpolarization can be observed, while during UP states cortical neurons are active (Steriade et al., 1993). Cortical DOWN and UP state dynamics can be objectively assessed by simultaneous LFP and single-unit activity recordings (Saleem et al., 2010). We analyzed the mean firing rate (z-scored) of single unit activity during the deactivated states to detect periods of silence (DOWN) and activity (UP) (**Figs. 3A, B**). Plotting of the LFP spectrogram over time revealed that the UP states were marked by pronounced power at frequencies >30 Hz in PV-Cre/NR1f/f mice, but not in PV-Cre mice (**Figs. 3A-C)**. In addition, compared to PV-Cre mice, the DOWN state in PV-Cre/NR1f/f was marked by increased power in a broader frequency range (including gamma frequencies; **Fig. 3A-C**). Decomposing the LFP signal using the Morlet wavelet transform also illustrates the alterations in mice lacking NMDAR activity in PV neurons (**Fig. 3D**). Detailed analysis of 4-second epochs in the LFP traces around the DOWN to UP state transitions (2 s before, and 2 s after state transition; **Figs. 3C, D**) revealed that loss of NMDAR in PV neurons was associated with significantly enhanced power specifically in the 30-60 Hz band (i.e., within the gamma range) and the 100-150 Hz (HFO) band, and significantly decreased power at slower frequencies (0.5-10 Hz) (**Figs. 3E-H**). Analysis of the transition-triggered average LFP revealed a less prominent DOWN state in PV-Cre/NR1f/f mice, compared to PV-Cre mice (**Fig. 3I)**, indicative of reduced hyperpolarization in the DOWN state of the slow oscillation (Saleem et al., 2010) in mice lacking NMDAR activity in PV neurons. The reduced hyperpolarization during DOWN states could contribute to the decreased power in lower LFP frequencies in PV-Cre/NR1f/f mice (**Fig. 3F**). Moreover, the increased gamma (30-60 Hz) and HFO power during state transitions (**Figs. 3G, H**) could contribute to the reduced phase-amplitude coupling during deactivated states in PV-Cre/NR1f/f mice (**Figs. 2D-H**). The altered LFP activities in PV-Cre/NR1f/f mice were associated with a variable DOWN state duration (**Figs. 3J, K**). However, PV-Cre/NR1f/f mice displayed significantly more DOWN and UP state events with shorter duration than PV-Cre mice (**Figs. 3J-M**), suggesting that the lack of NMDAR in PV neurons impacts the transition between DOWN and UP states.

**Figure 3.**
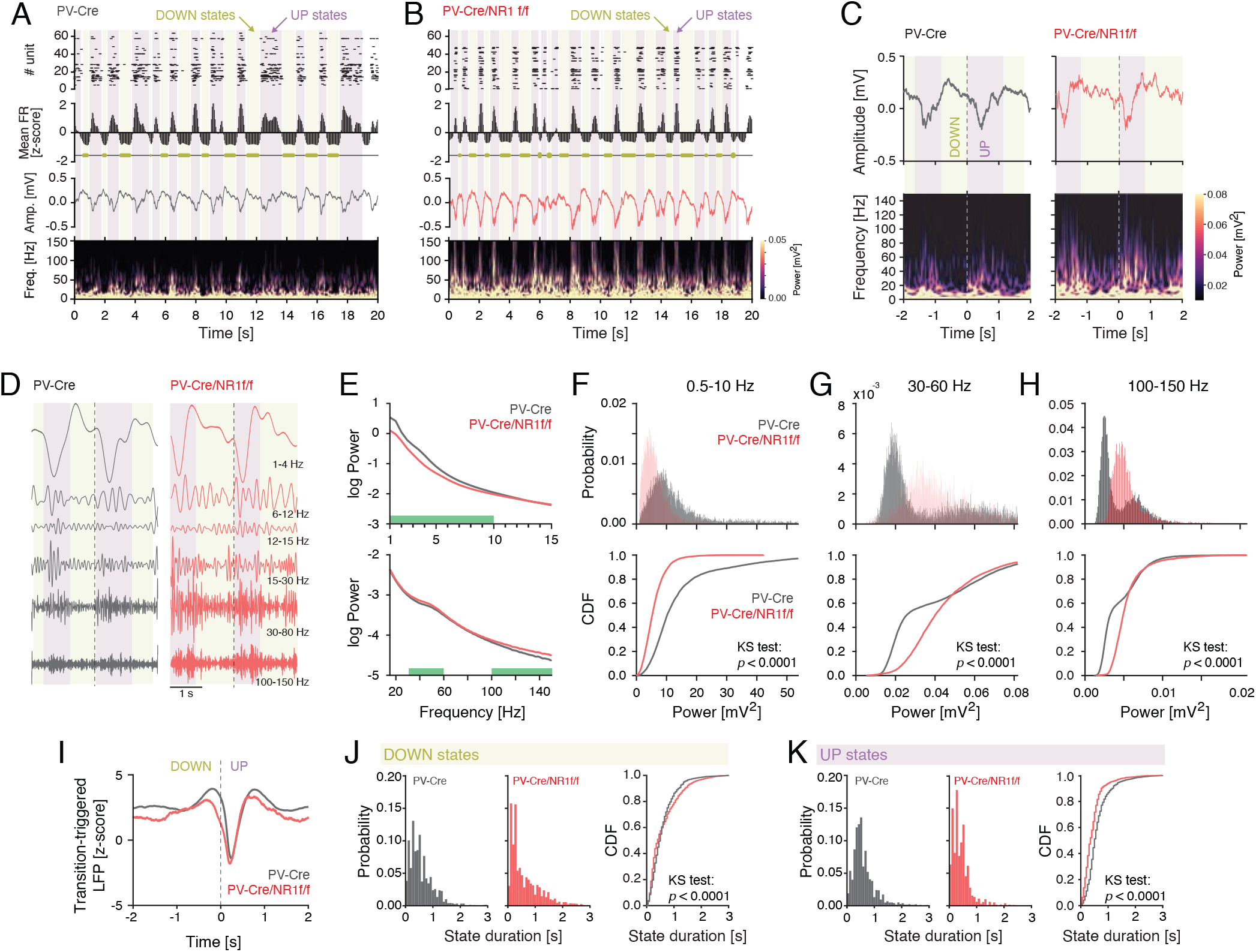
Mice with lack of NMDAR activity in PV neurons display altered LFP power during DOWN to UP state transitions and reduced duration of UP and DOWN states. **A**-**B**) Representative UP state (purple shading) and DOWN state (yellow shading) characteristics in a PV-Cre and a PV-Cre/NR1f/f mouse, respectively. From top to bottom: Raster plot of the recorded single units; the mean firing rate (FR, z-scored) with DOWN states indicated with yellow horizontal bars; the raw LFP; spectrogram (0-150 Hz) of the LFP. Mice with intact NMDAR activity in PV neurons (PV-Cre) display typical cyclic transitions between periods of very low/no spiking activity (DOWN states) and high spiking activity (UP states). Mice lacking NMDAR in PV neurons (PV-Cre/NR1f/f mice) display shorter DOWN and UP states than mice with intact NMDAR in PV neurons. In addition, the UP states have increased LFP amplitude, particularly in LFP frequencies >30 Hz, in PV-Cre/NR1f/f mice. **C)** Top: Representative unfiltered LFP traces (4 s) aligned to an example transition (time = 0 s) from a DOWN state (yellow shading) to a UP state (purple shading) in a PV-Cre and a PV-Cre/NR1f/f mouse, respectively. Bottom: Spectrograms (0-150 Hz) of the LFP traces in the top panels, highlighting the marked power increase in higher frequencies (>30 Hz) during UP states. Mice with lack of NMDAR activity display increased power in higher frequencies (> 30 Hz) during both DOWN and UP states. **D)** Representative Morlet-wavelets filtering of the LFP traces in (**C**) at different frequency bands. **E)** Comparison of the PSD of the LFP during the DOWN to UP state transition (−2 to 2 s) between PV-Cre vs PV-Cre/NR1f/f mice. Green bars: frequency bands with significant power difference between PV-Cre (n = 4; gray) and PV-Cre/NR1f/f mice (n = 3; red). **F-H)** Top: Probability histogram of power distribution in the frequency bands (**F**, 0.5-10 Hz; **G**, 30-60 Hz; **H**, 100-150 Hz) identified in (**E**) to hold differential power in PV-Cre vs PV-Cre/NR1f/f mice. Bottom: comparisons of the empirical cumulative distribution function of LFP power between PV-Cre and PV-Cre/NR1f/f mice. 0.5-10 Hz (*p* < 0.0001); 30-60 Hz (*p* < 0.0001); 100-150 Hz (*p* < 0.0001). PV-Cre (n = 4; gray) and PV-Cre/NR1f/f mice (n = 3; red). **I)** Mean transition-triggered LFP traces (4 s) of the DOWN to UP state transitions (time = 0 s) for PV-Cre (n = 4; gray) and PV-Cre/NR1f/f mice (n = 3; red). **J)** Probability histograms (left and middle) and empirical cumulative distribution function (right) of the duration of individual DOWN states in PV-Cre (n = 4; gray) and PV-Cre/NR1f/f mice (n = 3; red). Lack of NMDAR in PV neurons results in a significantly higher probability of shorter DOWN states, as confirmed by comparison of the empirical cumulative distribution function of the DOWN state duration between PV-Cre/NR1f/f and PV-Cre mice (*p* < 0.0001). **K)** Probability histograms (left and middle) and empirical cumulative distribution function (right) of the duration of individual UP states in PV-Cre (n = 4; gray) and PV-Cre/NR1f/f mice (n = 3; red). Lack of NMDAR in PV neurons results in a significantly higher probability of shorter UP states, as confirmed by comparison of the empirical cumulative distribution function of the UP state duration between PV-Cre/NR1f/f and PV-Cre mice (*p* < 0.0001). For the cumulative distribution functions, the Kolmogorov-Smirnov test was used to assess significance.

### Mice with lack of NMDAR activity in PV neurons display disorganized oscillatory activity during DOWN and UP states

As our results indicate increased 30-60 Hz gamma and HFO band power in PV-Cre/NR1f/f mice during DOWN and UP states (**Figs. 3E, G**), we next sought to clarify if this power increase was attributed to synchronous or asynchronous neuronal activity. Both highly synchronized neuronal activity and disorganized neuronal activity can increase the LFP power in different frequency bands, and power spectrum density (PSD) analysis is often used to determine the main oscillatory pattern emerging from the activity of a particular network in a specific moment. For instance, highly synchronized neuronal activity can lead to a stable build-up of LFP oscillations (i.e. a more sinusoidal-like oscillation), resulting in concentrated power in a narrow frequency band (e.g. sensory-evoked gamma oscillations), readily identified in the PSD. Disorganized neuronal activity generates a noisy and more irregular LFP, visible in the PSD as a more even power distribution over a broader frequency band (Uhlhaas et al., 2011). However, further analysis of the power spectrum is needed for quantitative comparison between different power distributions. To quantify the level of frequency dispersion (and consequently the level of synchronization in the neuronal activity) in the PSD, we performed analysis of the spectral entropy (Shannon Entropy) of the LFP (30-150 Hz), where a narrower power distribution results in lower entropy values, and a large power distribution results in higher entropy values, with no dependency on the LFP frequency (Valero et al., 2017).

Our analysis revealed significantly increased spectral entropy of the 30-150 Hz power spectrum during both UP and DOWN states in PV-Cre/NR1f/f mice compared to PV-Cre mice, indicating less synchronous LFP oscillations in mice lacking NMDAR activity in PV neurons (**Figs. 4A, C**). In addition to the level of synchronization, the rate of neuronal firing can modify the power spectrum of the LFP through spectral leakage, where high spiking activity results in broadband increases in LFP power (Manning et al., 2009; Ray and Maunsell, 2011; Scheffer-Teixeira et al., 2013). LFP frequencies higher than 200 Hz have been suggested to reflect the presence of substantial spectral leakage (Belluscio et al., 2012). Thus, to quantify a contribution of spectral leakage caused by increased spiking activity to the LFP power in the 30-150 Hz frequency band (broadband gamma + HFO bands, i.e., the aberrant bands in PV-Cre/NR1f/f mice), we calculated the ratio between the power of the 200-300 Hz band and the power of the 30-150 Hz band, hereafter referred to as the high-frequency index (HF index), for each DOWN and UP state event, respectively. This analysis demonstrated the occurrence of significantly more DOWN and UP states (events) with increased HF index in PV-Cre/NR1f/f mice than in PV-Cre mice (**Figs. 4B, D**), indicating a more substantial contribution of increased spiking activity to the 30-150 Hz power in the mutant mice. To understand how the observed less synchronous LFP oscillations (evident though significantly increased spectral entropy; **Figs. 4A, C**) in PV-Cre/NR1f/f mice related to the increased spiking activity (evident through significantly increased mean HF index; **Figs. 4B, D**), we explored the interaction between the spectral entropy values and the HF index. A large fraction of the events showed similar spectral entropy/HF index in the PV-Cre/NR1f/f and PV-Cre mice, however, PV-Cre/NR1f/f mice displayed a prominent occurrence of high entropy/high HF index events (**Figs. 4E, F**). This suggests that the increased power in the broadband gamma and HFO bands in the PV-Cre/NR1f/f mice is related to spectral leakage (measured by HF index) caused by increased firing rates associated with less synchronized LFP oscillations (measured by the spectral entropy).

**Figure 4.**
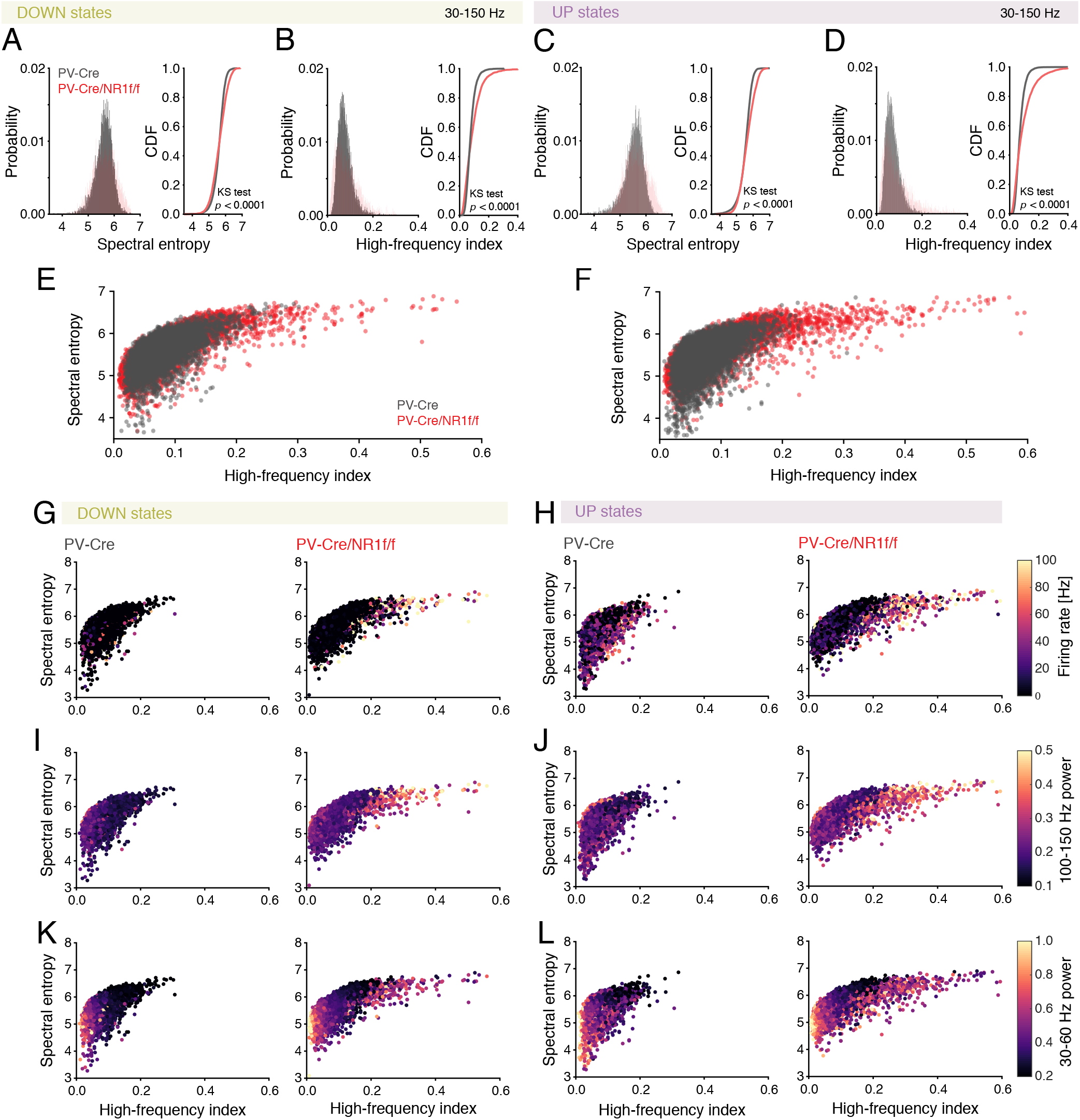
Mice with lack of NMDAR activity in PV neurons display asynchronous oscillatory activity associated with high firing rate and high-frequency (> 30 Hz) power during DOWN and UP states. **(A, B, E, G, I, K)**: DOWN states, **(C, D, F, H, J, L)**: UP states. **(A-L)**: PV-Cre (n = 4; gray) and PV-Cre/NR1f/f mice (n = 3; red). **A, C)** Right: Probability histogram of the spectral entropy of high LFP frequencies (30-100 Hz). Left: Comparison of the empirical cumulative distribution function of the spectral entropy demonstrates that PV-Cre/NR1f/f mice display significantly higher spectral entropy during both DOWN (**A**) and UP (**C**) states compared to PV-Cre mice (*p* < 0.0001). **B, D)** Right: Probability histogram of the HF index (calculated from the power ratio: 200-300 Hz / 30-150 Hz). Left: Comparison of the empirical cumulative distribution function of the HF index shows that PV-Cre/NR1f/f mice present significantly increased HF index during both DOWN (**B**) and UP (**D**) states compared to PV-Cre mice (*p* < 0.0001). **E-F)** Spectral entropy *versus* HF index. PV-Cre/NR1f/f mice, specifically, display a prominent occurrence of high entropy/high HF index events (PV-Cre: n = 9493 DOWN + UP events; PV-Cre/NR1f/f: n = 5971 DOWN + UP events). **G-L)** The high spectral entropy/high HF index events that are specific to mice with lack of NMDAR activity in PV neurons are associated with high firing rate and high-frequency (> 30 Hz) power. Projection (as colormaps) of the firing rate (**G, H)**, power of 100-150 Hz LFP (**I, J)** and power of 30-60 Hz LFP (**K, L**) over spectral entropy *versus* HF index (PV-Cre: n = 9493; PV-Cre/NR1f/f: n = 5971 DOWN + UP events). **G-H**) The high spectral entropy/high HF index events in PV-Cre/NR1f/f mice are marked by high firing rate. **I-J)** The high spectral entropy/high HF index events in PV-Cre/NR1f/f mice are marked by high 100– 150 Hz LFP power. **K-L)** Low spectral entropy/low HF index in both PV-Cre and PV-Cre/NR1f/f mice are marked by high 30-60 Hz (within gamma range) power. However, PV-Cre/NR1f/f mice in addition display high spectral entropy/high HF index events marked by high 30-60 Hz LFP power. For the cumulative distribution functions, the Kolmogorov-Smirnov test was used to assess significance.

To directly delineate the contributions of single-unit activity and activity in the frequency bands with differential power in PV-Cre/NR1f/f and PV-Cre mice (**Fig. 3E**) to the occurrence of high spectral entropy/high HF index events, we projected the average firing rate, the 30-60 Hz power, and 100-150 Hz power, respectively, against the spectral entropy and the HF index. This identified that the high spectral entropy/high HF index events specific to the PV-Cre/NR1f/f mice were associated with increased firing rates during both DOWN and UP states (**Figs. 4G, H**), showing that the less synchronous LFP oscillations identified by increased spectral entropy in PV-Cre/NR1f/f mice (**Figs. 4A, C**) are associated with increased firing rates. The high spectral entropy/high HF index events in PV-Cre/NR1f/f mice were furthermore marked by increased power in the HFO (100-150 Hz) band (**Figs. 4I, J**), i.e., higher firing rates and higher HFO power co-existed within events, suggesting that increased power in the HFO band is related to increased neuronal firing and to less synchronous LFP oscillations. Importantly, increased power in the 30-60 Hz band (i.e., within the gamma range) was both in PV-Cre mice and PV-Cre/NR1f/f mice associated with low spectral entropy/low HF index events during the DOWN and UP states (**Figs. 4K, L**). However, in PV-Cre/NR1f/f mice, increased power in the 30-60 Hz band was in addition associated with high spectral entropy/high HF index events (**Figs. 4K, L**). The presence of this dual distribution of increased 30-60 Hz band power in PV-Cre/NR1f/f mice, but not in PV-Cre mice, suggests that a proportion of the DOWN and UP events in PV-Cre/NR1f/f mice contain gamma activity (30-60 Hz) with the same properties as gamma in PV-Cre mice (i.e., lower spectral entropy and lower HF index). However, the high spectral entropy/high HF index in other events with increased 30-60 Hz power in PV-Cre/NR1f/f mice suggests an origin from spectral leakage caused by increased spiking activity. Together our analyses suggest that increased neuronal activity contributes to the increased power in the 30-60 Hz and 100-150 Hz bands observed specifically in high entropy/high HF index DOWN and UP state events in mice lacking NMDAR activity in PV neurons.

### Temporal dynamics of mPFC PV interneurons and excitatory neurons during DOWN to UP state transition

Both cortical PV interneurons and excitatory neurons fire preferentially during the UP state (Puig et al., 2008; Tahvildari et al., 2012) but PV interneurons also display low firing activity during the DOWN state, which has been suggested to influence DOWN-to-UP transition probability (Zucca et al., 2017). As our LFP analyses indicated increased neuronal firing and altered state transitions in mice lacking NMDAR activity in PV neurons, our next investigations focused on single unit activities during DOWN to UP transitions. As before (Kim et al., 2016, 2020), we used opto-tagging to identify the activity of PV interneurons in PV-Cre and PV-Cre/NR1f/f mice injected with AAV-DIO-ChR2-mCherry (**Figs. S1A-C**). Application of blue light resulted in significantly increased firing in a set of neurons (**Fig. S2A-C**). Prefrontal inhibitory PV interneurons provide potent inhibition onto pyramidal neurons in the local network (Kim et al., 2016), and in agreement with this the application of blue light resulted in silencing of neurons in the local circuit (**Fig. S2D**). In contrast, blue light application did not modulate prefrontal activities in PV-Cre mice injected with AAV-DIO-eYFP (n = 4; **Figs. S2E-G**). The spike waveforms of the light-activated units (n = 5; **Figs. S2H-K**) were signified by short half-valley width and low peak-to-valley amplitude ratio, typical of prefrontal narrow-spiking (NS) interneurons (**Fig. S3A;** (Kim et al., 2016). These typical features were further used to classify recorded units into neuronal subtypes by calculation of the NS probability in a Gaussian mixture model (GMM) clustering (**Fig. S3B**). Based on the NS probability units were classified as wide-spiking (WS) putative pyramidal neurons (n = 155, NS probability < 0.3) or NS putative interneurons (n = 63, NS probability > 0.9), respectively. Unclassified units (n = 98, NS probability between 0.3 and 0.9) were not further analyzed (**Figs. S3C, D**). A second GMM clustering based on peak distribution (calculated with normalized amplitude waveform) was applied to further classify all NS units (n = 63 units) as putative PV (n = 14 units, probability > 0.9) and NS (n = 49 units, probability < 0.3). This classification was further confirmed by the calculation of the spike waveform similarity index with light-induced spike waveforms, where the PV interneurons within the NS population presented a waveform similarity index >95%. (**Figs. S3E, F**). In summary, our analyses identified WS, NS, and putative PV interneurons, respectively, with distinct spike waveform features (**Fig. S3G**).

Isolation of 1-second windows around the DOWN to UP state transitions (0.5 seconds before, and 0.5 seconds after state transition) and analysis of the mean firing rate (z-scored) of individual mPFC WS, NS, and PV interneurons, respectively, revealed that all three classes of neurons displayed firing primarily in the UP state in both PV-Cre/NR1f/f and PV-Cre mice **(Figs. 5A-I**). The average firing rate of WS neurons was significantly increased in PV-Cre/NR1f/f1 mice during DOWN states, compared to PV-Cre mice (**Figs. 5A, B**). Further, while the mean WS firing rate during UP states was similar between PV-Cre/NR1f/f and PV-Cre mice (**Fig. 5B**), the firing, as well as the firing onset, was more temporally dispersed in PV-Cre/NR1f/f mice compared to PV-Cre mice (**Figs. 5A-C**), suggesting reduced precision in the coordination of WS activities in mice lacking NMDAR activity in PV neurons. We found this aberration to be accompanied by a significantly increased peak latency of the PV firing during the UP states in PV-Cre/NR1f/f mice (**Fig. 5F**). In addition, plotting of the peak-width at half-height (PWHH) of the mean firing curve of individual PV interneurons indicated that the firing of PV interneurons in PV-Cre/NR1f/f mice was more temporally variable across the UP states, compared to PV-Cre mice (**Fig. 5F**). Overall the data shows that lack of NMDAR activity led to reduced precision in the spike timing of mPFC PV interneurons and consequently, impaired WS spike precision, consistent with previous findings in the somatosensory cortex (Carlén et al., 2012). The dynamics of NS putative interneurons during state transition was similar in PV-Cre/NR1f/f and PV-Cre mice (**Figs. 5G-I**). Altogether, the analyses of single-unit activities support the view that cortical PV interneurons are integral to the neuronal dynamics in the mPFC during state transitions (Massi et al., 2012; Zucca et al., 2017).

**Figure 5.**
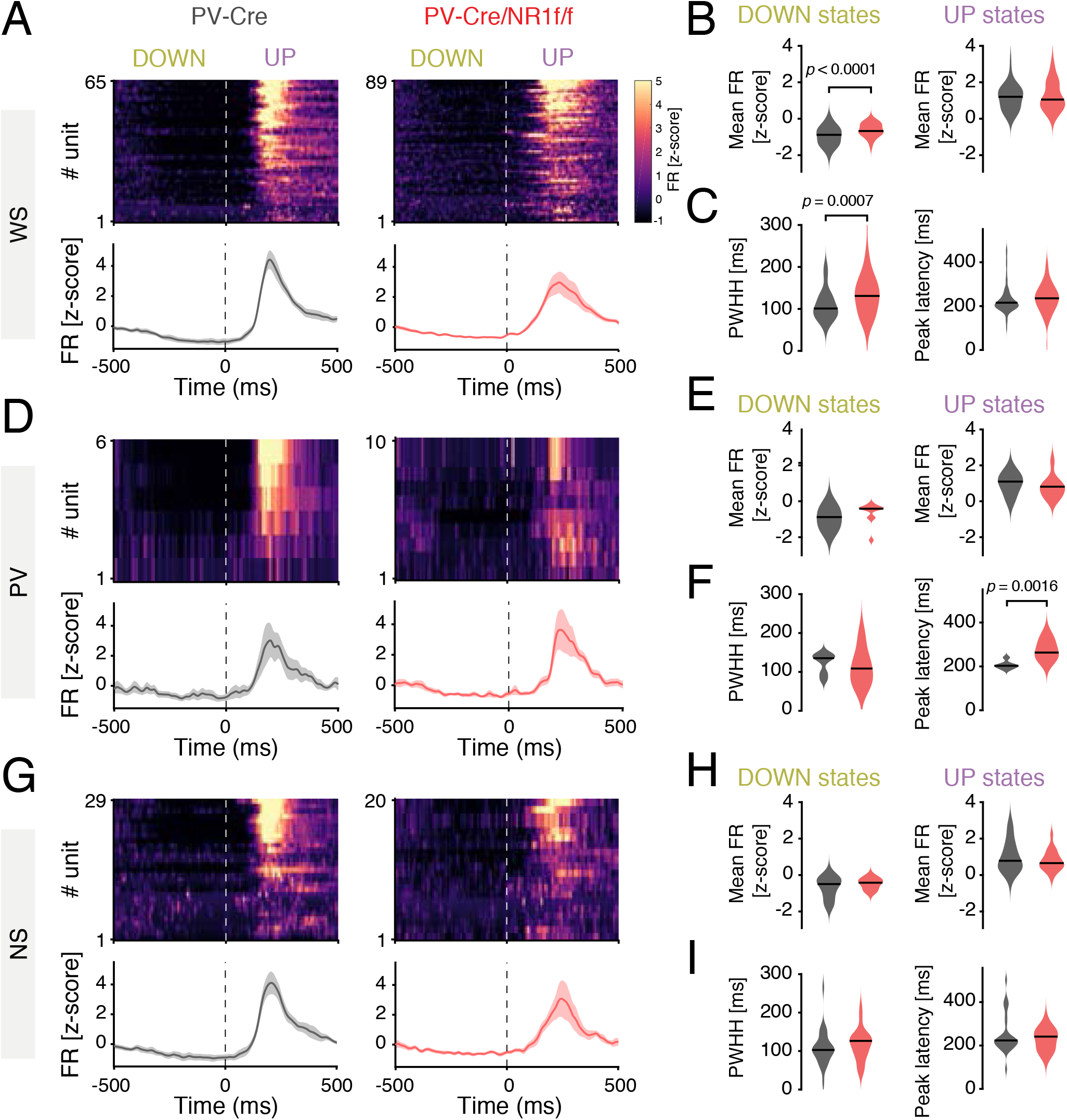
Mice with lack of NMDAR activity in PV neurons display altered temporal dynamics of prefrontal cortex single-unit activity during DOWN to UP state transitions. **A)** Firing rate (FR) dynamics of WS neurons during DOWN-to-UP state transition (−500 to 500 ms, 0 = transition (dashed line). PV-Cre (gray; n = 65 units) and PV-Cre/NR1f/f mice (red; n = 89 units). Top: Peri-stimulus time histograms showing mean FR (z-score) of single WS units. Units are sorted according to their mean FR (z-score) during the UP state (0 to 500 ms). Bottom: Mean FR (z-score) for all WS units. **B-C)** Firing properties of WS neurons during DOWN to UP state transition (−500 to 500 ms). **B)** The WS FR (z-score) is significantly higher in PV-Cre/NR1f/f mice than in PV-Cre mice during DOWN states. DOWN: PV-Cre: −0.95; PV-Cre/NR1f/f: −0.74; *p* <0.0001; UP: PV-Cre: 1.17; PV-Cre/NR1f/f: 1.01; *p* = 0.4802. **C)** Left: violin plots of the peak width at half-height (PWHH) of the average FR of individual WS neurons showing significantly increased spike time variability of WS in PV-Cre/NR1f/f mice compared to PV-Cre mice. Right: Peak latency of the spiking of individual WS neurons. PWHH: PV-Cre: 100.9 ms; PV-Cre/NR1f/f: 130.9 ms; *p* = 0.0007; peak latency: PV-Cre: 215 ms; PV-Cre/NR1f/f: 235 ms; *p* = 0.0661. **D-F)** Same as (**A-C**) but for putative PV interneurons. PV-Cre (gray; n = 6 units) and PV-Cre/NR1f/f mice (red; n = 10 units). **(E)** The PV FR (z-score) during DOWN (left) and UP states (right). DOWN: PV-Cre: −0.87; PV-Cre/NR1f/f: −0.41; *p* = 0.2198; UP: PV-Cre: 1.10; PV-Cre/NR1f/f: 0.81; *p* = 0.3132. **(F)** Left: violin plots of the peak width at half-height (PWHH) of the average FR of individual PV interneurons. Right: The peak latency of the spiking of individual PV interneurons is significantly increased in PV-Cre/NR1f/f mice compared to PV-Cre mice. PWHH: PV-Cre: 133.2 ms; PV-Cre/NR1f/f: 106.3 ms; *p* = 0.3132; peak latency: PV-Cre: 202.5 ms; PV-Cre/NR1f/f: 262.5 ms; *p* = 0.0016. **G-I)** Same as (**A-C**) but for NS putative interneurons. PV-Cre (gray; n = 29 units) and PV-Cre/NR1f/f mice (red; n = 20 units). **H)** The NS FR (z-score) during DOWN (left) and UP states (right). DOWN: PV-Cre: −0.55; PV-Cre/NR1f/f: −0.48; *p* = 0.1532; UP: PV-Cre: 0.74; PV-Cre/NR1f/f: 0.61; *p* = 0.2944. I) Left: violin plots of the peak width at half-height (PWHH) of the average FR of individual NS interneurons. Right: Peak latency of the spiking of individual NS interneurons. PWHH: PV-Cre: 102.8 ms; PV-Cre/NR1f/f: 126.5 ms; *p* = 0.3581; peak latency: PV-Cre: 220 ms; PV-Cre/NR1f/f: 237 ms; *p* = 0.7052. For (**A, D, G**) data shown as mean ± SEM (shaded area). For violin plots, black lines represent median. Two-tailed unpaired t-test was used to assess significance if data passed the D’Agostino & Pearson normality test, if not, the Mann Whitney test was used.

### Distinct characteristics of broadband gamma oscillations induced by ketamine

Local mPFC administration of NMDAR antagonists is known to elicit marked alterations of oscillatory patterns, inducing a state with lower LFP amplitude and increased power in faster frequencies (> 30 Hz) (Kulikova et al., 2012; Lee et al., 2017). We questioned if the ketamine-induced oscillatory alterations are related to asynchronous neuronal activity (e.g., increased firing rate and/or decreased spike precision that we found to be associated with increased high-frequency power in PV-Cre/NR1f/f mice). For this, the silicon probe was moved to a new mPFC recording site and 30 min baseline single-unit and LFP oscillations were recorded. S(+)-ketamine was thereafter locally applied into the mPFC (**Methods**) and the effects recorded for 30 min (60 min recording time in total; PV-Cre, n = 3 mice; PV-Cre/NR1f/f; n = 3 mice). In accordance with earlier reports (Kulikova et al., 2012), ketamine application triggered a rapid shift in the LFP oscillations in PV-Cre mice, from predominantly low frequencies of high amplitude to a state with persistent high frequency (>30 Hz) activity of low amplitude (**Figs. 6A-C**). The LFP oscillations in the ketamine induced state were desynchronized, as revealed by plotting of the autocorrelogram of the LFP over time (60 min) (**Fig. 6D**). In contrast, ketamine application did not overly affect the LFP oscillations in mice lacking NMDAR activity in PV neurons (**Figs. 6A-D)**. Instead, PV-Cre/NR1f/f mice displayed low frequency activity of high amplitude both before and after application of ketamine (**Figs. 6A, B**). Ketamine increased the power of most frequency bands in PV-Cre mice (**Fig. 6C**), and the power change was significantly larger than in PV-Cre/NR1f/f mice (**Fig. 6C**). Consequently, PV-Cre mice displayed significantly higher broadband gamma and HFO power after local application of ketamine compared to PV-Cre/NR1f/f mice (**Figs. 6E-G**). In addition, the low-delta (0.5-1.5 Hz) power was significantly lower in PV-Cre mice than in PV-Cre/NR1f/f mice (**Figs. 6E-G**). Our results confirm that NMDAR in PV neurons are a central target of NMDAR antagonists (Homayoun and Moghaddam, 2007; Hakami et al., 2009; Carlén et al., 2012; Cohen et al., 2015; Jadi et al., 2016; Picard et al., 2019; Hudson et al., 2020), and show that PV interneurons are central to ketamine’s effects on activities in prefrontal networks.

**Figure 6.**
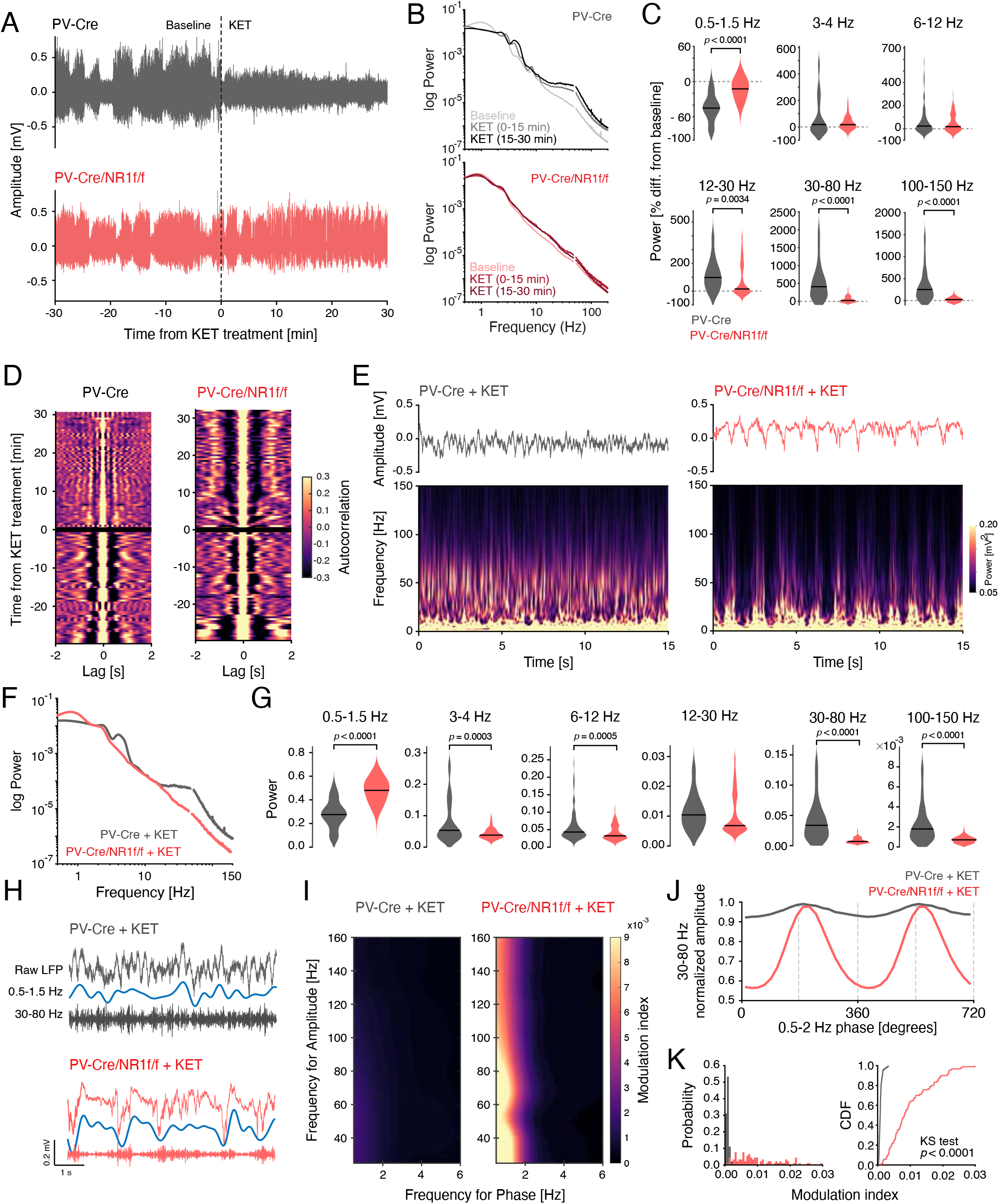
Distinct spectral characteristics of the LFP induced by ketamine. **A)** Representative unfiltered mPFC LFP traces (60 min) from a PV-Cre (top) and a PV-Cre/NR1f/f (bottom) mouse recorded under urethane anesthesia. Ketamine (KET) was locally applied at t = 0; dashed line. In PV-Cre mice, ketamine application switched the LFP oscillations from predominantly low frequencies of high amplitude to higher frequencies of low amplitude. In contrast, ketamine application did not cause any major changes in the mPFC LFP oscillations in PV-Cre/NR1f/f mice. **B)** Mean PSD of the LFP for the PV-Cre mice (gray; n = 3) and PV-Cre/NR1f/f mice (red; n = 3) during baseline (−30-0 min), 0-15 min, and 15-30 min after ketamine application. **C)** Power change (from baseline) of different frequency bands induced by ketamine application (−30-0 vs 0-30 min). Power of low-delta band decreased significantly more in PV-Cre mice (gray; n = 3) than in PV-Cre/NR1f/f mice (red; n = 3) after ketamine application while the beta band, broadband gamma and HFO power were significantly more increased in PV-Cre than PV-Cre/NR1f/f mice. Low-delta: PV-Cre: −45.22 %; PV-Cre/NR1f/f: −12.49 %; *p* <0.0001; high-delta: PV-Cre: +18.40 %; PV-Cre/NR1f/f: +17.10 %; *p* = 0.8428; theta: PV-Cre: +22.5 %; PV-Cre/NR1f/f: +17.20 %; *p* = 0.4450; beta: PV-Cre: +98.3 %; PV-Cre/NR1f/f: +15 %; *p* = 0.0034; broadband gamma: PV-Cre: +408.2 %; PV-Cre/NR1f/f: +24 %; *p* <0.0001; HFO: PV-Cre: +251.4 %; PV-Cre/NR1f/f: +21.2 %; *p* <0.0001). **D)** Evolution of the autocorrelograms of the LFP traces in (**A**) through time (60 min), before (−30-0 min) and after (0-30 min) ketamine application (t = 0; black line). Colorbar represents the LFP autocorrelation coefficients. Ketamine lowers the LFP autocorrelation in PV-Cre mice, indicating desynchronized LFP oscillations, while the LFP in PV-Cre/NR1f/f mice at large does not change. **E-G)** Direct comparison of LFP activities after ketamine application in PV-Cre mice (gray; n = 3) and PV-Cre/NR1f/f mice (red; n = 3). **E)** Top: 15 s of the unfiltered LFP traces in (**A**) after ketamine application (0-30 min). Bottom: Spectrograms (0-150 Hz) of the LFP traces in the top panels, colorbar represents LFP power. Ketamine application triggered a state with persistent high frequency (>30 Hz) activity of low amplitude in the LFP oscillations in PV-Cre mice, while PV-Cre/NR1f/f mice displayed low frequency activity of high amplitude. **F)** Mean PSD (0-150 Hz) of the LFP after ketamine application (0-30 min), demonstrating decreased power in lower frequencies and increased power in higher frequencies in PV-Cre, but not PV-Cre/NR1f/f, mice. **G)** Power of the different frequency bands after ketamine application (0-30 min). PV-Cre mice display significantly higher power in most frequency bands compared to PV-Cre/NR1f/f mice but significantly lower power in the low-delta band. Low-delta: PV-Cre: 0.27; PV-Cre/NR1f/f: 0.48; *p* <0.0001; high-delta: PV-Cre: 0.053; PV-Cre/NR1f/f: 0.038; *p* = 0.0003; theta: PV-Cre: 0.043; PV-Cre/NR1f/f: 0.032; *p* = 0.0005; beta: PV-Cre: 0.0104; PV-Cre/NR1f/f: 0.0068; *p* = 0.0579; broadband gamma: PV-Cre: 0.0336; PV-Cre/NR1f/f: 0.0068; *p* <0.0001; HFO: PV-Cre: 0.0018; PV-Cre/NR1f/f: 0.0007; *p* <0.0001. **H-K)** Phase amplitude co-modulation after ketamine application (0-30 min). **H)** Representative unfiltered LFP traces (top), filtered low-delta (0.5-1.5 Hz; middle), and broadband gamma (30-80 Hz; bottom) bands under ketamine in a PV-Cre (gray) and a PV-Cre/NR1f/f (red) mouse. **I)** Representative phase-amplitude comodulograms of a PV-Cre (left) and a PV-Cre/NR1f/f mouse (right) after ketamine application (0-30 min), colorbar represents the modulation index (MI). Ketamine decreases the modulation of broadband gamma amplitude by the low-delta phase in PV-Cre, but not in PV-Cre/NR1f/f, mice. **J)** Mean broadband gamma amplitude (normalized) at different phases of the low-delta cycles for PV-Cre mice (n = 3; gray) and PV-Cre/NR1f/f mice (n = 3; red) after ketamine application (0-30 min). **K)** Left: Distribution of the MI probability between low-delta phase and broadband gamma amplitude in PV-Cre mice (n = 3) and PV-Cre/NR1f/f mice (n = 3) after ketamine application (0-30 min). Right: Comparison of the empirical cumulative distribution function of MI shows that PV-Cre/NR1f/f mice have significantly more 30 s epochs of higher MI than PV-Cre mice (*p* < 0.0001). In (**C, G**), black line: median. Two-tailed unpaired t-test was used to assess significance if data passed the D’Agostino & Pearson normality test, if not, the Mann Whitney test was used. For the cumulative distribution function, the Kolmogorov-Smirnov test was used to assess significance.

We, furthermore, questioned how ketamine-induced broadband gamma oscillations are coordinated by lower LFP frequencies. Based on our previous results, we focused on the coupling between the low-delta (0.5-2 Hz) phase and the broadband gamma (30-80 Hz) amplitude, which we found to be significantly lower in PV-Cre/NR1f/f mice than in PV-Cre mice during deactivated states, a reflection of desynchronized LFP oscillations in PV-Cre/NR1f/f mice (**Figs. 2C-F**). Interestingly, the delta-gamma phase-amplitude coupling was significantly decreased in PV-Cre mice compared to PV-Cre/NR1f/f mice after ketamine application (**Figs. 6H-K)**. The altered response to ketamine in mice lacking NMDAR activity in PV neurons was also evident in the single-unit dynamics. While the ketamine state was characterized by a more tonic firing in PV-Cre mice, the single-unit activity in PV-Cre/NR1f/f mice after ketamine application remained oscillating between silent (DOWN) and active (UP) periods (**Figs. 7A, B)**, with the mean firing rate of both WS neurons and NS interneurons (the latter including putative PV interneurons) being significantly higher in PV-Cre mice than in PV-Cre/NR1f/f mice (**Figs. 7C, D**). Together, the analyses show that local ketamine application induces desynchronized LFP oscillations, increased firing rates, and increased power in a (very) broad range of LFP oscillations in mice with intact NMDAR activity in PV neurons. In addition, the coupling between broadband gamma amplitude and the phase of low-frequency oscillations decreases.

**Figure 7.**
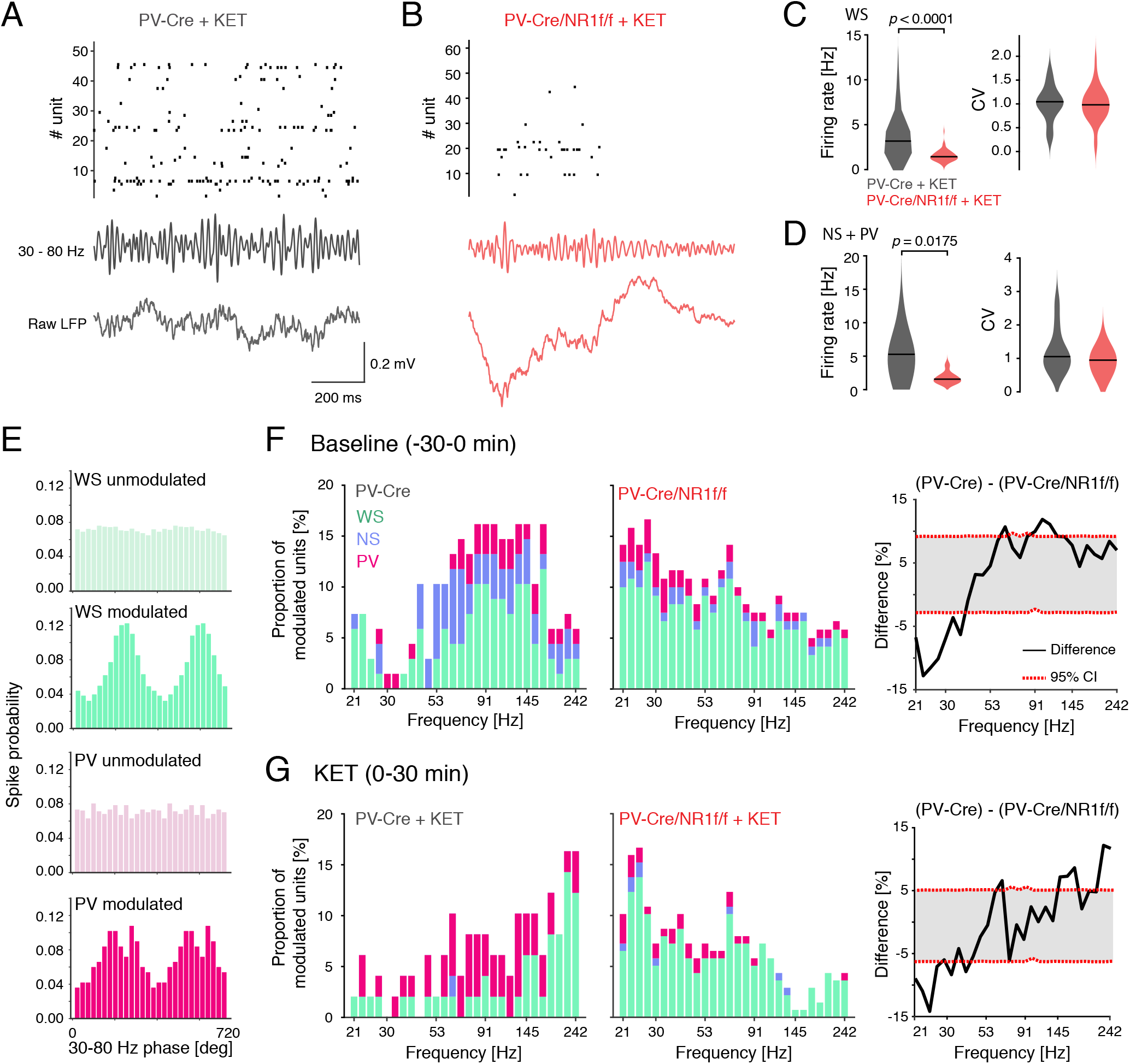
Firing patterns and spike-LFP entrainment in the mPFC during baseline, and after ketamine application. **A-B)** Representative (1 s) single-unit activity (top), filtered LFP in the 30-80 Hz band (middle) and raw LFP traces (bottom) for a PV-Cre (**A**) and a PV-Cre/NR1f/f mouse (**B**) after ketamine application. **C)** FR properties of WS neurons (PV-Cre: n = 17 units; PV-Cre/NR1f/f: n = 69 units) after ketamine application (0-30 min). Left: the FR of WS neurons is significantly higher in PV-Cre mice than in PV-Cre/NR1f/f mice after ketamine application. Right: The coefficient of variation (CV) of the inter-spike interval does not differ between PV-Cre (n = 3; gray) and PV-Cre/NR1f/f (n = 3; red) mice. FR: PV-Cre: 3.28; PV-Cre/NR1f/f: 1.52; *p* < 0.0001; CV: PV-Cre: 1.03; PV-Cre/NR1f/f: 0.97; *p* = 0.6751. **D)** FR properties of NS+PV putative interneurons (PV-Cre: n = 10 units; PV-Cre/NR1f/f: n = 17 units) after ketamine application (0-30 min). Left: The FR of NS+PV interneurons is significantly higher in PV-Cre mice than in PV-Cre/NR1f/f mice after ketamine application. Right: The CV of the inter-spike interval does not differ between PV-Cre (n = 3; gray) and PV-Cre/NR1f/f (n = 3; red) mice. FR: PV-Cre: 5.27; PV-Cre/NR1f/f: 1.56; *p* = 0.0175; CV: PV-Cre: 1.05; PV-Cre/NR1f/f: 0.95; *p* = 0.3093. **E)** Histograms of the spike modulation by broadband gamma (30-80 Hz) phase. Greens; representative single WS neurons, pinks; representative single PV interneurons. **F)** Spike-LFP entrainment during baseline (−30-0 min). The proportion of neurons significantly (*p* < 0.01) modulated by different LFP frequencies (21-242 Hz; log scale) in PV-Cre mice (n = 3; left) and PV-Cre/NR1f/f (n = 3, middle) mice. WS (green): PV-Cre: n = 22 units; PV-Cre/NR1f/f: n = 90 units; NS (blue): PV-Cre: n = 7 units; PV-Cre/NR1f/f: n = 12 units); PV (pink): PV-Cre: n = 3 units; PV-Cre/NR1f/f: n = 5 units. The spiking in PV-Cre mice is predominantly modulated by higher frequencies, and in PV-Cre/NR1f/f mice by lower frequencies. Right: Difference in the proportion of neurons (WS + NS + PV) significantly (*p* < 0.01) modulated by different LFP frequencies (21-242 Hz) between PV-Cre (n = 3) and PV-Cre/NR1f/f (n = 3) mice. The modulation of spiking by higher LFP frequencies is significantly stronger in PV-Cre than in PV-Cre/NR1f/f mice. **G)** Spike-LFP entrainment of the neurons in (**F**) after ketamine application (0-30 min). Ketamine shifts the modulation of spiking in PV-Cre mice (left) to higher LFP frequencies (>200 Hz) while the spiking in PV-Cre/NR1f/f mice (middle) at large is not affected. Right: after ketamine application the modulation of frequencies >200 Hz is significantly stronger in PV-Cre, than in PV-Cre/Nr1f/f, mice. For (**C** and **D**) black line: median. Two-tailed unpaired t-test was used to assess significance if data passed the D’Agostino & Pearson normality test, if not, the Mann Whitney test was used.

### Prefrontal spike-LFP entrainment during baseline and after ketamine application

LFP oscillations contribute to spatiotemporal coordination of neuronal activity, modulating the spiking probability according to the phase and frequency of the field oscillations (Watson et al., 2018). As we found (increased) spiking in PV-Cre/NR1f/f mice contaminating the LFP power > 30 Hz (**Figs. 4G-L**), we as a next step analyzed the spike-LFP coupling for different frequencies, with a specific focus on the broadband gamma (30-80 Hz) and HFO bands. Using normal circular distribution statistics (see **Methods**), we first calculated the modulation of each neuronal population (WS, and NS, PV, respectively) to the higher frequencies of the LFP (21-242 Hz) during baseline (**Fig. 7E-G**). To measure the strength of population entrainment to the different LFP frequencies, we calculated the proportion neurons statistically (*p* <0.01) modulated by each specific LFP frequency. During baseline (0-30 min before ketamine application) a large proportion of the mPFC WS neurons in the PV-Cre mice was entrained by higher frequencies (∼91-145 Hz; **Fig. 7F)**. The PV interneurons in PV-Cre mice showed similar patterns of modulation (**Fig. 7F**). A large proportion of the WS neurons in PV-Cre/NR1f/f mice were in contrast entrained by a broader, and lower, frequency spectrum (21-90 Hz; **Fig. 7F**). A bootstrap procedure (see **Methods**), used to detect significant differences between PV-Cre/NR1f/f and PV-Cre mice in the entrainment of spiking to the different LFP frequencies, identified significantly stronger spike coupling to high frequency LFP oscillations (∼90-120 Hz) in PV-Cre mice than in PV-Cre/NR1f/f mice (**Fig. 7F**). In contrast, the spiking in PV-Cre/NR1f/f mice displayed significantly stronger modulation by <40 Hz oscillations than in PV-Cre mice (**Fig. 7F**).

We next investigated how the desynchronization induced by ketamine alters spike-LFP coupling in the mPFC network, and the plausible dependence on NMDAR activity in PV neurons. This revealed that ketamine remarkably reorganized the entrainment of neurons to the LFP in mice with intact NMDAR activity in PV neurons. Specifically, the preferential modulation of WS neurons shifted from ∼90 Hz towards faster LFP frequencies (>200 Hz) (**Fig. 7G**), while PV interneurons were modulated by a broader range of LFP frequencies during the ketamine state than during baseline (**Fig. 7G**). In contrast, the altered spike-LFP entrainment in PV-Cre/NR1f/f mice seen during baseline were at large not affected by the application of ketamine (**Fig. 7G**), strongly indicating that PV interneurons (with intact NMDAR activity) are integral to proper spike-LFP coupling in the mPFC. The differential spike-LFP entrainment associated with ketamine-induced broadband gamma vs the increased spontaneous (baseline) broadband gamma in mice lacking NMDAR activity in PV neurons, suggest separate circuit underpinnings (see further in Discussion).

## Discussion

There is an emerging view that (spontaneous) broadband gamma oscillations that are measured via a power increase across a wide frequency range at baseline or task-free conditions may be a reflection of increases in the overall level of circuit activity rather than of rhythmic neural activity that is synchronized across neurons (see Sohal and Rubenstein, 2019). While feedback inhibition by cortical PV interneurons is central to rhythmic circuit activity underlying more narrow-band increases in gamma power, the role in broadband gamma increases is currently unclear. However, given the important role of PV interneurons in cortical excitation/inhibition balance, reduced PV interneurons function is likely to increase the overall levels of circuit activity, which could cause a broadband power increase across a wide range of frequencies. We here show that mice lacking NMDAR activity in PV neurons display increased power across a (very) broad frequency band, including the broadband gamma band (30-80 Hz) and the HFO band (100-150 Hz). Importantly, we find the increased high-frequency power to be associated with LFP oscillations and neuronal activity with decreased synchronicity. The main findings distinguishing asynchrony in PV-Cre/NR1f/f mice were i) reduced coupling between low-delta phase and broadband gamma amplitude of the LFP, ii) significantly increased spectral entropy in the 30-150 Hz power spectrum (reflecting a broader LFP power distribution and less synchronous LFP oscillations), and iii) a significantly higher HF index (reflecting a significantly larger contribution of spiking activity to the 30-150 Hz power by spectral leakage), compared to mice with intact NMDAR activity in PV neurons. As mentioned earlier, a broad LFP power distribution is considered to reflect a noisy and more irregular LFP caused by disorganized neuronal activity (Uhlhaas et al., 2011), and our single-unit analysis confirmed significantly increased spike latency of PV interneurons and significantly increased variability in the spike-timing of excitatory neurons in mice lacking NMDAR activity in PV neurons (Carlén et al., 2012). Earlier work has demonstrated that LFP activity >80 Hz can reflect either true neuronal oscillations or spectral leakage of spiking activity (spike “contamination”) (Ray and Maunsell, 2011; Scheffer-Teixeira et al., 2013). We show here that the high spectral entropy/high HF index events in mice lacking NMDAR activity in PV neurons hold spike contamination caused by increased firing rates, leading to increased power in the 30-60 Hz and HFO bands. However, the presence of also low spectral entropy/low HF index events with increased power in the 30-60 Hz band suggests the coexistence of oscillations within the gamma band caused by spectral leakage and genuine LFP oscillations in PV-Cre/NR1f/f mice. The reduced precision in the spike timing of mPFC PV interneurons could contribute to the spectral leakage of spiking activity associated with the broadband increase of LFP power in PV-Cre/NR1f/f mice. It is notable that the asynchronies in PV-Cre/NR1f/f mice primarily manifested during the deactivated state, a cortical state marked by synchronous activity (Steriade, 2006; Harris and Thiele, 2011).

PV interneurons have been implicated in the regulation of cortical DOWN and UP state transitions (Zucca et al., 2017) and we show here that the increased temporal variability of PV firing in PV-Cre/NR1f/f mice is accompanied by increased firing rates of WS neurons during DOWN states - a state normally marked by hyperpolarization. Optogenetic inhibition of cortical PV interneurons has been shown to trigger a DOWN-to-UP transition, which has led to the suggestion that PV firing (although weak) during DOWN state is fundamental to maintain the cortical circuitry in the silent state (Zucca et al., 2017). In line with this, we found the deactivated state to be fragmented in mice lacking NMDAR activity in PV neurons, with significantly reduced DOWN and UP state duration (plus increased variability of the DOWN state duration in PV-Cre/NR1f/f mice compared to PV-Cre mice). The reduced hyperpolarization during DOWN state could directly contribute to the decreased delta power, and increased power in frequencies >30 Hz, apparent during the deactivated states in mice lacking NMDAR activity in PV neurons. NMDAR hypofunction in PV neurons could give rise to increased excitatory activity through different mechanisms. Reduced postsynaptic NMDAR activity in PV neurons could reduce the PV excitability, and hence the PV activity (Korotkova et al., 2010; Carlén et al., 2012). More recent findings suggest that presynaptic activity of NMDARs in PV interneurons enhances the probability of GABA release, including in the PFC (Pafundo et al., 2018). Reduced presynaptic NMDAR activity would reduce the inhibitory postsynaptic currents (IPSCs) and the strength of PV-to-pyramidal inhibition (Jadi et al., 2016; Pafundo et al., 2018). As we here delete the ubiquitous NR1 subunit, both pre- and postsynaptic changes in NMDAR activity could plausibly contribute to the observed alterations.

Local ketamine application resulted in a desynchronized state, characterized by lower amplitude and higher frequencies of LFP oscillations, and reduced modulation of gamma amplitude by slow-delta phase in mice with intact NMDAR activity in PV neurons. Interestingly, systemic ketamine administration has been shown to alter sleep-like oscillations (induced by pentobarbital) in the frontoparietal cortex in similar ways, and the associated LFP changes found to be related to ketamine-induced alterations in the thalamic reticular nucleus (TRN) and thalamocortical networks (Mahdavi et al., 2020). Thus, our results on reduced synchronization of LFP activity might be similar to the effect of systemic ketamine of thalamic networks (Mahdavi et al., 2020), given that local application of ketamine affects connected structures (Kulikova et al., 2012). In agreement, local blockade of NMDAR activity by application of MK-801 has been demonstrated to propagate through connected neural circuits (Lee et al., 2017). Previous *in vivo* studies have demonstrated that NMDAR antagonists, including ketamine, increase the firing of prefrontal neurons (Wood et al., 2012; Molina et al., 2014). We show here that local ketamine application increases the firing of both prefrontal WS neurons and NS interneurons. The observation that the power of cortical high frequencies LFP oscillations (50-180 Hz) and the firing rates in the local circuits positively correlate with spike-LFP entrainment in the 50-180 Hz band suggests that local neuronal activity underlies the generation of high frequency LFP oscillations (Watson et al., 2018). Analysis of spike-LFP entrainment, thus, can offer insight to mechanisms underlying high frequency LFP oscillations. The baseline spike-LFP entrainment in mice with intact NMDAR activity in PV neurons were predominantly observed in the 50-180 Hz band, with local ketamine application shifting the entrainment to higher frequencies (>200 Hz). However, mPFC neurons in mice lacking NMDAR activity in PV neurons were entrained by LFP oscillations of a broader frequency range (and particularly by lower frequencies), and ketamine did not change the spike-LFP coupling. The differential spike-LFP entrainment associated with ketamine-induced broadband gamma vs the increased spontaneous (baseline) broadband gamma in mice lacking NMDAR activity in PV neurons, suggest separate circuit underpinnings. Acute (NMDAR antagonists), and long-term (genetic), PV hypofunction, respectively, clearly entail distinct circuit alterations, the latter including not only activity alterations but also structural changes. Dendritic and axonal structural changes in PV interneuron can give rise to lasting disorganization of local network activities (del Pino et al., 2013; Hamm et al., 2017; Guyon et al., 2020), alteration not replicated by ketamine-induced suppression of PV inhibition (Hamm et al., 2017).

Sleep and circadian rhythm dysfunctions are comorbidities of schizophrenia. It is interesting to notice that the fragmentation of the deactivated state in the current study parallels findings of fragmentation specifically during NREM sleep in mouse models of schizophrenia involving PV interneuron dysfunction (Phillips et al., 2012). Cortical states transitions under sleep-wake cycles (reviewed in (Brown and McKenna, 2015) and under urethane anesthesia (Clement et al., 2008), involve complex interactions between cortical and subcortical networks, with notable involvement of cholinergic (Clement et al., 2008; Kim et al., 2015; Tikhonova et al., 2018), and long-range GABA-ergic PV neurons in the basal forebrain (McKenna et al., 2020; McNally et al., 2020), and GABA-ergic PV neurons in the TRN (Steriade, 2006; Cohen et al., 2015; Mahdavi et al., 2020). As an example, optogenetic stimulation of PV neurons in the basal forebrain shortens the NREM sleep states, without affecting REM states (McKenna et al., 2020). The global knock-out of NMDAR in PV neurons in PV-Cre/NR1f/f mice could affect long-range PV neurons in the basal forebrain and their function in coordination of cortical brain states, possibly contributing to the observed fragmentation of the deactivated states. Clearly, the lack of NMDAR in prefrontal PV interneurons could, in addition, affect the local processing of inputs from subcortical regions regulating arousal and cortical states (Bogart and O’Donnell, 2018; Ährlund-Richter et al., 2019). In summary, our investigations demonstrate that broadband power increases in gamma oscillations can be a result of asynchronous neuronal activity, and that the power increases can arise by different circuit mechanisms. Our results, thus, give support for the view that increases in circuit activity resulting from increased excitation/inhibition ratio can be reflected by increased power of broadband gamma oscillations in the LFP. Importantly, our findings demonstrate that PV interneurons do not only contribute to genuine narrow-band gamma oscillations that depend on rhythmic feedback inhibition (Cardin et al., 2009; Buzsáki and Wang, 2012), but can also underlie power increases in broadband gamma oscillations associated with cognitive dysfunction.

## Methods

### Animals

All procedures were performed in accordance and compliance with the guidelines of the Stockholm Municipal Committee for animal experiments and the Karolinska Institutet in Sweden (approval N 43/14). To generate mice lacking NMDAR specifically in PV interneurons (PV-Cre-NR1f/f mice), PV-Cre knock-in mice (JAX stock #008069, RRID:IMSR_JAX:008069) were crossed to mice carrying the ‘floxed’ NR1 alleles (JAX stock #005246, RRID:IMSR_JAX:005246), as previously characterized (Carlén et al., 2012). Anesthetized electrophysiology recordings were performed on adult (15-20 weeks of age) PV-Cre/NR1f/f (n = 3 females, n = 1 male) and PV-Cre mice (n = 6 females, n = 2 males). PV-Cre/NR1f/f and PV-Cre mice were bred at the Karolinska Institute, Sweden. Animals were housed in groups (3-5 mice/cage) using individually ventilated caging systems (Sealsafe plus, GM 500, Tecniplast, Buggugiate, Italy), under standardized conditions with a 12-hour light-dark cycle (light 7:00 am), stable temperature (20 ± 1°C) and humidity (40-50%) with access to food (R70 Standard Diet, Lactamin AB, Vadstena, Sweden; 4.5 gm% fat, 14.5 gm% protein, 60 gm% carbohydrates) and water ad libitum.

### Viral injection

The injections were performed in a biosafety cabinet class 2 with sterile surgical environment. A micropipette (10 µl graduated borosilicate glass capillary; Wiretrol I Calibrated Micropipettes, Drummond, Broomall, PA, U.S.A.) prefilled with mineral oil was fixed into the Quintessential Stereotaxic Injector (Stoelting, Wood Dale, IL, U.S.A.) and filled with 0.5 µL of virus (titer ∼ 10^12^) dyed with *Fast Green* 2.5% (Electron Microscopy Sciences, Hatfield, PA, U.S.A.). The mice (∼8 weeks old) were injected subcutaneously with the analgesic buprenorphine (0.1 mg/kg; Temgesic, Indivior Europe Limited, Dublin, Ireland) 30 min prior to surgery. The mice were placed on a heating blanket (37°C) in the stereotaxic apparatus (Harvard Apparatus, Holliston, MA, U.S.A.) and anesthetized with 2% isoflurane in oxygen, and the eyes were covered with eye lubricant (Viscotears, Novartis, France). The mice’s reflexes were tested and the isoflurane level was reduced to 1 or 1.5% over the course of the surgery. The scalp was shaved and lidocaine (4 mg/kg) was injected subcutaneously before skin incision. The connective tissue around the skull was gently removed as needed for clear viewing of the Bregma. The craniotomy was performed unilaterally above the mPFC: AP: 1.8 mm anterior to Bregma, ML: 0.3 mm lateral to midline. Sterile saline 0.9 % was regularly applied over the skull while drilling to remove bone dust and control heat generation. The total diameter of the burr hole was rarely larger than the diameter of the drill bit (around 100 µm). Using a syringe needle the dura was carefully opened, and the micropipette tip was lowered until reaching the target location (prelimbic area (PL) of the mPFC; AP: +1.8 mm; ML: +0.3 mm; DV: −1.4 mm, relative to the surface). 0.5 µL of AAV-DIO-ChR2-mCherry (rAAV2/Ef1A-DIO-hChR2(H134R)-mCherry; 4 × 10e12 viral particles/ml, from UNC Vector Core, NC, U.S.A.) was injected at a rate of 0.1 µL/min. Mice used for control of optogenetic artifacts (PV-Cre; n = 4) were injected with AAV-DIO-eYFP (rAAV5/EF1a-DIO-eYFP; 4 × 10e12 particles/ml; UNC Vector Core, NC, U.S.A.). The micropipette was retained at the injection site for 10 minutes after injection and before retraction. The edges of the skin were pulled together and sealed with small amounts of cyanoacrylate glue (Vetbond Tissue Adhesive, Henry Schein, Melville, NY, USA), and carprofen (5 mg/kg; Norocarp, Norbrook Laboratories, Ireland) was given subcutaneously. The mice were monitored until completely recovered from the anesthesia. An additional dose of carprofen was administered 24 h after surgery.

### Electrophysiological recordings, optogenetic stimulation, and S(+)-ketamine injection

Five to seven weeks after the virus injection, the mice were anesthetized with urethane (1.1 g/Kg in sterile saline 0.9%, i.p.) and fixed in a stereotaxic apparatus (Digital Stereotaxic with Dual Manipulators; Leica Microsystems, Nussloch GmbH, Germany) on a heating blanket (Harvard Apparatus) to maintain the body temperature at 37 ± 0.5°C and kept under 0.25% isoflurane anesthesia for the course of the surgery. Using the above-described coordinates, we made a new craniotomy over the initial burr hole, large enough to allow positioning of an optical fiber (details below) and a silicon probe (A4×2-tet-5mm-150-200-312-A32 Neuronexus, Ann Arbor, MI, USA) coupled to a 32 channel headstage preamplifier (Neuralynx, Bozeman, MT, USA) liked to a Digital Lynx 4SX acquisition systems (Neuralynx). Small amounts of saline 0.9% were continuously dropped on the craniotomy in order to prevent brain surface dryness. An additional craniotomy was made over the parietal cortex to fix a ground-screw, used as a reference. Then, we carefully positioned the silicon probe in the target position (AP: 1.8 mm anterior to Bregma, ML: 0.3 mm relative to midline, DV: −2 mm). We monitored in real-time the electrophysiological signal for 30 min to allow the brain tissue to recover and optimize the recording condition. Two different recordings were performed. First, 50 min of recording was performed to monitor different brain states induced by urethane. Second, a 60 min recording was performed to assess prefrontal oscillatory and neuronal activity alterations induced by S(+)-ketamine. In this new recording session, after moving the silicon probe (DV: ∼0.2 mm), single-unit and LFP oscillations were monitored for 30 min before and after S(+)-ketamine (100 µg in sodium chloride 0.9% solution) applied over the craniotomy where the silicon probe was implanted. One PV-Cre mouse was not included in this part of the analysis due to poor recording quality. The recordings were acquired using the Cheetah 5.0 acquisition and experiment control software (Neuralynx). Electrical signals were divided, pre-amplified (1000x) and filtered for local field potentials (LFPs) and single-unit activity. The biological signal was sampled at 32 kHz and band-pass filtered between 600–6000 Hz to record spikes and between 0.5–500 Hz to record LFPs.

For optogenetic manipulations, we positioned an unjacketed optical fiber (200 µm core diameter; numerical aperture: 0.22; ThorLabs, Newton, NJ, USA) 1.5 mm above the recording site using a tilted insertion (10 to 15 degrees of slope; 0.3 mm below the brain surface), in order to attenuate photoelectric artifacts by reducing the contact exposure to the light (Cardin et al., 2010). Optical stimulation was generated by a 473 nm laser source (100 mW; CNI Lasers, Changchun, China) triggered through a data acquisition board (BNC-2110, National Instruments, Austin, TX, USA) controlled by a custom-made LabView (National Instruments) program. A broad range of light-stimulation frequencies (8, 16, 24, 32, 40, 48, 60, 80, and 90 Hz) was applied randomly in bouts of 3 s of 1-ms pulses width with 5 mW. Laser power was measured at the tip of the optical fiber before each experiment by a PM100D power meter console (ThorLabs).

### Tissue processing, immunostaining and imaging

The mice were deeply anesthetized with pentobarbital (60 mg/ml; Apotek Produktion & Laboratorier AB, Sweden) and transcardially perfused with 1x phosphate-buffered saline (PBS, 0.1 M, pH 7.4) followed by 4% paraformaldehyde (PFA) in PBS (0.1 M, pH 7.4). The perfused brain was removed from the skull and postfixed with 4% PFA in 0.1 M PBS at 4°C for 16 h. The brain was thoroughly washed in 0.1 M PBS and thereafter sectioned (40 µm thickness) using a vibratome (Leica VT1000, Leica Microsystems). The sections were stored in 1x PBS at 4°C.

Vibratome cut (free floating) tissues were permeabilized for 1 h with 1x TBST (0.3% Triton-X in 1x TBS), blocked for 1 h at room temperature (RT) with 10% normal donkey serum in 1x TBST, and thereafter incubated with primary antibodies (rabbit polyclonal anti-PV; Cat# PV25, RRID:AB_10000344, Swant, Switzerland; at a dilution of 1:1000) in 1x TBST at RT for 12-24 h. The sections were thereafter washed three times in 1x TBST, and incubated for 3-5 h with a secondary antibody (Alexa Fluor® 488 anti rabbit; Cat# 711-545-152, RRID:AB_2313584, Jackson ImmunoResearch, Cambridge, United Kingdom; at a dilution of 1:500) in 1x TBST. The sections were thereafter consecutively washed with 1x TBST, 1x TBS and 1x PBS (10 min each). All sections were mounted on glass slides (Superfrost Plus, Thermo ScientificTM, Waltham, MA, U.S.A.) and coverslipped using 50:50 Glycerol:1x PBS mixed with DAPI (added for visualization of nuclei).

Images were acquired at 10 or 20x magnification with a fluorescent microscope (Leica DM6000B) connected to a Hamamatsu Orca-FLASH 4.0 C11440 digital camera (Hamamatsu, Hamamatsu City, Japan). Images were spatially registered to the Allen Mouse Brain Common Coordinate Framework version 3 (ABA_v3) reference atlas (Lein et al., 2007) using the QuickNII tool (RRID:SCR_016854; (Puchades et al., 2019) and SBA Composer (Bakker et al., 2015).

### Data analysis

All data analysis was conducted using custom scripts written in MATLAB (The MathWorks, Natick, MA, USA) and GraphPad Prism version 8.00 (GraphPad Software, La Jolla, CA, USA).

### Brain state classification

LFP oscillations were divided in 30 s epochs. Using a Gaussian mixture model, epochs of the LFP were classified in deactivated, activated and transition (unclassified) states based on the power of the 0.5-2 Hz and 3-45 Hz frequency bands for an initial clustering. The classified epochs were manually refined by eliminating epochs at the cluster edge. Spectrum content of each state was analyzed to confirm proper classification. Transition epochs were discarded from the analysis.

### UP state and DOWN state classification

DOWN and UP states during deactivated states were detected based on the z-scored average firing rate for each mouse, with individually tuned thresholds. Firing rates were calculated with a 0.01 s bin and a Gaussian window of 0.4 s. Units with an average firing rate <0.01 spike/s were not included. Analyses were conducted only with paired events, where paired events correspond to the most recent DOWN state around an UP state event. UP state events with no surrounding DOWN state were excluded.

### Spectral analysis

Power Spectral Density (PSD) were estimated using Welch’s method (MATLAB algorithm), with 1 s Hamming window, with 50% overlap and a 5120 Fast-Fourier transform (FFT) points. Normalized power was obtained by dividing the PSD estimation by the integrated power over all frequencies (0.5-200 Hz). For visualization of LFP oscillations in specific frequency bands, a band-pass filter using two-way least-squares FIR filtering (*eegfilt* function from EEGLAB toolbox) was applied (Delorme and Makeig, 2004).

Morlet-wavelet spectrograms of the LFP oscillations were calculated using a wavelet convolution, built with a Morlet wavelet with frequency resolution of 0.2 Hz and frequency bounds of 0.5 and 150 Hz. Each frequency of the resultant spectrum was normalized by the sum of power spectrum from 0.5 to 200 Hz. The power distribution over the frequency bands was calculated by the extraction of the amplitude envelope (Hilbert transform) from the wavelet-decomposed signal in order to obtain representative Morlet-wavelets of different frequency bands (1-4 Hz, 6-12 Hz, 12-15 Hz, 15-30 Hz, 30-80 Hz, 100-150 Hz). The transition-triggered average field potential was calculated as the mean LFP during 4 s for all events, centered at the DOWN to UP state transition.

### Phase-amplitude modulation index

Phase-amplitude coupling was estimated by the modulation index (MI) as described in Tort et al., 2010. MI is a normalized measure that reflects how well the amplitude of a faster oscillation is phase-locked to an underlying lower cycle. Comodulation maps were constructed calculating the MI at phase frequencies from 0.5 Hz to 6 Hz (in steps of 1 Hz), and amplitude frequencies from 30 Hz to 150 Hz (in steps of 5 Hz). Shannon entropy of the distribution of mean amplitudes per phase (divided into 18 bins) in each frequency bin was calculated to obtain the cross-frequency MI for each 30 s epoch. Specifically, the MI for low-delta-broadband gamma, and low-delta-HFO were calculated and, then, averaged for group comparisons.

### Spectral Entropy and High-Frequency Index

To quantify the level of frequency dispersion in the PSD, we performed analysis of the spectral entropy of the LFP (30-150 Hz). Based on previous work using this measure (Valero et al., 2017), we determined the spectral entropy of the LFP by calculating the Shannon Entropy, given by Eq. (1).

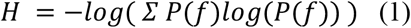

Where P(f) is the normalized power spectrum calculated in 35 bins of frequency from 30 to 150 Hz band.

To quantify the level of spectral leakage by spiking activity contributing to the power of the LFP, we determined the high-frequency index (referred as HF index; with values from 0 to 1). We first calculated the power at 30-150 Hz and 200-300 Hz frequency bands by extracting the amplitude envelope (Hilbert transform) from the wavelet-decomposed signal, and then divided the power at 200-300 Hz frequency band by the power at 30-150 Hz.

### Local Field Potential autocorrelogram

To exemplify how ketamine changes the level of synchronicity of prefrontal oscillations, LFP autocorrelograms were calculated for all 30 s epochs. For this, we measured the similarity for each 30 s LFP and their respective shifted lags (from −2 to 2 seconds), using the function *xcorr* from Matlab with *coeff* normalization, thus obtaining cross-correlation arrays for each epoch. Then, we built a two-dimensional plot with the cross-correlation values for all epochs along time (60 min), indicating the autocorrelation values as a colormap.

### Spike sorting

Spikes were manually sorted into individual units (putative neurons) off-line based on analysis of peak amplitude, valley amplitude, peak-valley ratio, spike width, waveform energy and principal components using the MClust 4.4 offline spike sorting software (A.D. Redish; available at http://redishlab.neuroscience.umn.edu/MClust/MClust.html). Cluster quality was evaluated by the isolation distance and L-ratio (Schmitzer-Torbert et al., 2005). Clusters with isolation distance <12 or L-ratio >0.3 were excluded. Autocorrelation and interspike interval (ISI) histograms charts were inspected for all putative neurons. In cases in which the autocorrelation showed absolute refractory period violations (spike counts at periods <1.2 ms), we improved cluster separation, otherwise, the cluster was excluded.

### Units classification

The units were classified into wide-spiking (WS) putative pyramidal neurons and narrow-spiking (NS) putative interneurons based on the distribution of (1) the peak-to-valley ratio (the ratio between the amplitude of the initial peak and the following trough) and (2) the half-valley width of each spike waveform. For the objective classification of units between WS and NS, a Gaussian mixture model (GMM) was fit to the unit’s features, as previously described in Kim et al., 2016. Units with low classification confidence (*p* >0.05) were discarded. To identify putative PV interneurons, the NS population was further classified based on the combination of waveform similarity index and the magnitude of the repolarization (spike waveform peak) (**Figs. S3D, E**). The waveform similarity index (*r*) was calculated by correlating each spike waveform with the mean spike waveform of light-activated units. The value of 1 indicates identical spike waveforms. NS neurons with a peak amplitude (normalized) <0.8 and high waveform similarity index (>95%) were classified as PV interneurons.

### Single-units activity analyses

To determine single-unit activity during DOWN to UP state transitions, peri-event time histograms (10 ms bins, 500 ms pre-event, 500 ms post-event) were calculated and convoluted with a Gaussian Kernel of 50 ms and normalized using z-score. For each unit, we calculated the mean firing rate at DOWN and UP states, the latency to the peak (maximum firing rate; relative to the DOWN to UP transition), and the peak width at half height (PWHH), which reflects the level of spike timing precision around the peak. Units with firing rate <0.1 spike/s were excluded for these analyses.

To determine single-unit activity under ketamine, we calculated for each 30 s epoch interspike intervals (ISI) histograms for every unit to obtain the mean firing rate (FR = 1/ISI) and the coefficient of variation (CV), as the ratio of standard deviation of ISI by the average ISI. (CV = std(ISI)/mean(ISI)). To assess the effects of optogenetic stimulation on the activation of mPFC neurons, neuronal activity relative to the light events was expressed in a peri-event time histogram (PETH) and a spike raster (rastergram) for each neuron. To dismiss any light artifacts in light-responsive units, the spike waveform shape similarity was determined by calculating the correlation coefficient (*r*) between two average spike waveforms (1 s before and after the light onset) for each unit.

### Measurement of spike-LFP entrainment using von Mises statistics

We quantified the entrainment of spikes to the LFP as previously described in Schomburg et al., 2014. In brief, we extracted the entrainment of spikes to the LFP by filtering the LFP with a second-order Butterworth filter and extracting the phase using the Hilbert transform. By calculating the phases of the filtered-LFP where spikes occurred, we tested whether the spike was significantly modulated by the LFP using a von Mises distribution (or circular normal distribution) statistics. The entrainment of spikes by the LFP was calculated on a log scale from 21 to 242 Hz and the mean length of the resultant vector was taken for each frequency and plotted in color code. To avoid spurious spike entrainment to high frequencies due to LFP contamination by the spike itself, we used the LFP on the shank next to the shank where the specific cell was recorded (200 µm distance). Units recorded from electrodes with noisy LFP signal were excluded. Units included in the analysis: WS units (PV-Cre: n = 22; PV-Cre/NR1f/f: n = 90), NS units (PV-Cre: n = 7; PV-Cre/NR1f/f: n = 12), Putative PV units (PV-Cre: n = 3; PV-Cre/NR1f/f: n = 5).

A bootstrap procedure was used for identification of significant differences in the spike-LFP entrainment between PV-Cre and PV-Cre-NR1f/f mice (during baseline and in response to ketamine application, respectively). The spike-LFP entrainment for the two mouse cohorts was calculated in randomly shuffled LFP frequencies, and the proportion of neurons modulated for each frequency was recalculated. This procedure was repeated for 10000 times and a confidence interval of 95% was built. The 95% CI was then used to identify the frequencies in which the difference between both cohorts are significant.

### Statistical Analysis

Normal distribution was evaluated in all data sets using the D’Agostino & Pearson normality test. If the data fulfilled the criteria for normal distribution, we analyzed the data using two-tailed unpaired or paired t-test. If the data did not pass the normality test, we used the Mann Whitney test. An analysis of variance (Two-way ANOVA), and Bonferroni post-hoc comparisons in case of significant interactions, was used to compare the data in **Figs. 1E-G**. For cumulative distribution functions, the Kolmogorov-Smirnov test was used. Only values of *p* < 0.05 were considered statistically significant.

## Author contributions

N.G., J.P.L., C.L.A. and M.C. conceived the experiments. N.G. and C.L.A. conducted *in vivo* electrophysiology experiments. N.G., L.R.Z., E.F.O., H.K. and C.L.A. analyzed electrophysiology data. N.G. performed histological analysis. N.G., E.F.O. and C.L.A. prepared figures. N.G., L.R.Z, E.F.O. J.P.L., C.L.A. and M.C. wrote the manuscript. All authors reviewed the manuscript.

## Acknowledgments

This study was financed in part by a STINT Program Joint Brazilian-Swedish Research Collaboration grant, a CAPES-STINT program grant (n° 99999.009883/2014-02) and by the Coordenação de Aperfeiçoamento de Pessoal de Nível Superior – Brasil (CAPES) – Finance Code 001. C.L.A. was supported by a São Paulo Research Foundation (FAPESP) grant (n° 2012/07107-2). M.C. was supported by a Wallenberg Academy Fellow in medicine grant (n° KAW 2012.0131) from the Knut and Alice Wallenberg Foundation, by a European Research Council Starting Grant (n° 337069), by the Swedish Research Council (n° 2016-02700) and Karolinska Institutet (n° 2016-00139). We would like to thank Flávio A. G. Mourão for help with data analysis, and Stephanie Rogers and Adriano Tort for the valuable comments on this manuscript.

---

**Figure S1.**
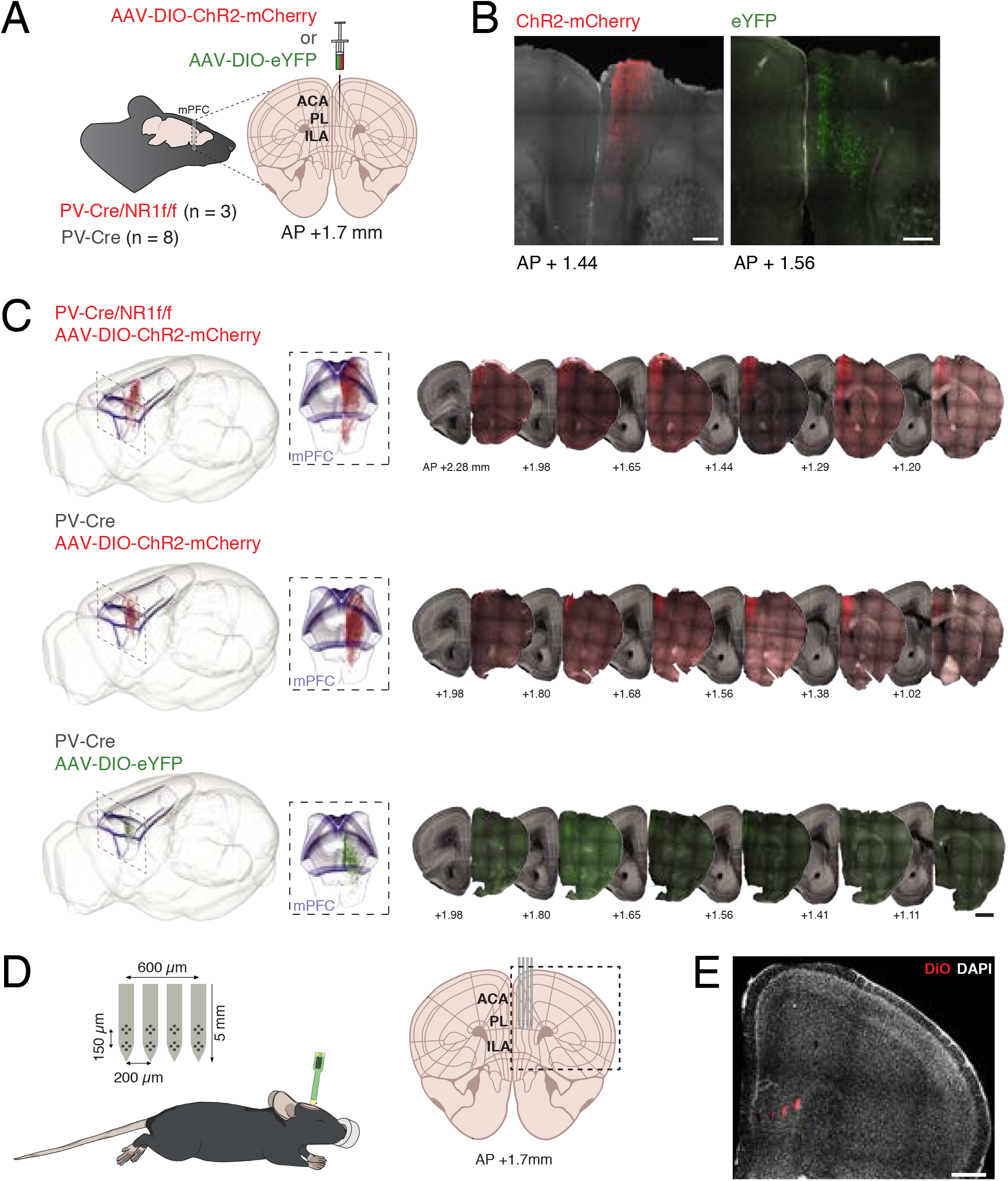
Virus expression and positioning of silicon probes. **A)** Illustration of viral injections. For optogenetic targeting of mPFC PV interneurons, AAV5-DIO-ChR2-mCherry (red) was unilaterally injected into the mPFC of PV-Cre/NR1f/f (n = 3) and PV-Cre (n = 4) mice. AAV5-DIO-eYFP (green) was unilaterally injected into the mPFC of PV-Cre (n = 4) for control of optogenetic artifacts (**Figs. S2E-G**). **B)** Coronal sections of the mPFC showing expression of ChR2-mCherry (red; left) in a representative PV-Cre/NR1f/f mouse, and eYFP (green; right) in a representative PV-Cre mouse. **C)** Left: Representative 3D illustration of detected mPFC expression of ChR2-mCherry (top: PV-Cre/NR1f/f mouse; middle: PV-Cre mouse) and eYFP (bottom; PV-Cre mouse) registered to the Allen Mouse Brain Common Coordinate Framework version 3 (ABA_v3) reference atlas. Box; the mPFC (ACA, ILA, PL according to ABA_v3). Middle: close up (front view) of the box in left. Right: Representative coronal sections used for detection and 3D plotting of virus expressing neurons. **D)** Illustration of the experimental setup. Electrophysiological recordings were conducted using a silicon probe in urethane-anesthetized mice. The four-shank eight-tetrodes silicon probe was targeted to the PL area in the mPFC. **E)** Example DiO (red) labelling of the four shank tracts of a silicon probe. Nuclei are counterstained by DAPI (white). ACA: Anterior Cingulate are; PL: Prelimbic area; IL: Infralimbic area; AP: antero-posterior axis, from Bregma. Scale bars: (**B**): 300 µm; (**C**): 1 mm; (**E**): 500 µm. Brain silhouette in (**A**) and mouse in (**D**) were sourced from https://scidraw.io/ and adapted from (Kennedy, 2020; Tyler and Kravitz, 2020) respectively.

**Figure S2.**
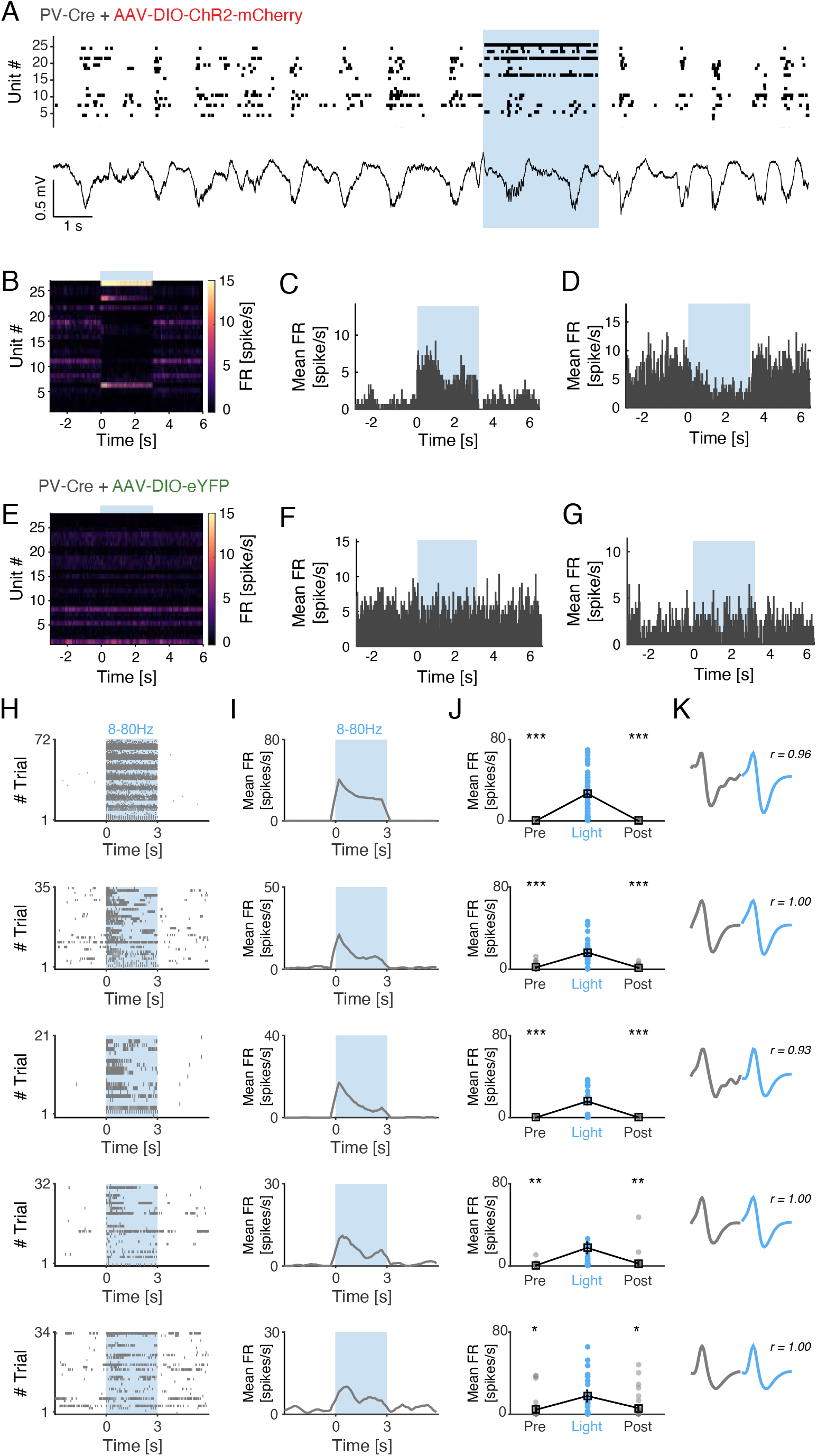
Opto-tagging of ChR2-expressing mPFC PV interneurons. **A)** Example spiking activity of single-units (top; n = 27) and raw LFP (bottom) recorded (20 s) in the mPFC of a PV-Cre mouse injected with AAV5-DIO-ChR2-mCherry. The spiking of recorded units was modulated in different ways by application of 3 s blue light (blue shading; 473 nm, 46 mW/mm2, 3 ms pulses, 40 Hz). **B)** Mean FR of the neurons in (**A**) across 90 trials. **C)** Peri-stimulus time-histogram demonstrating increased FR of a light-activated unit in (**A**) in response to blue light application. **D)** Peri-stimulus time-histogram demonstrating decreased FR of a mPFC unit in (**A**) in response to light activation of local PV interneurons. **E-G)** Blue light application did not modulate the firing rates in PV-Cre mice injected with AAV-DIO-eYFP. Experimental settings as in (**A**). **G)** Mean FR of mPFC neurons (n = 28) across 90 trials in an example PV-Cre mouse injected with AAV-DIO-eYFP. **E-F**) No light artifacts were detected as shown in these two unmodulated units. **H)** Spiking activity of the five light-responsive mPFC PV interneurons (PV-Cre + PV-Cre/NR1f/f mice) in response to application of 3 s blue light (blue shading; 473 nm, 46 mW/mm2, 3 ms pulses). Light frequencies used: 8, 16, 24, 32, 40, 48, 80 Hz. Only trials with mean FR >0.1 spike/s during baseline (t = -3-0 s) are included. **I-J**) Mean FR of the 5 light-responsive units in (**H**). All five units showed significant increased spiking in response to blue light application. **K**) Average spike waveform of the five light-responsive units in (**H**). Spontaneous (gray) and light-evoked (blue) spike waveforms exhibit very high similarity. r = Waveform similarity. Two-tailed paired t-test was used to assess significance. *p < 0.05, **p < 0.01, ***p < 0.001.

**Figure S3.**
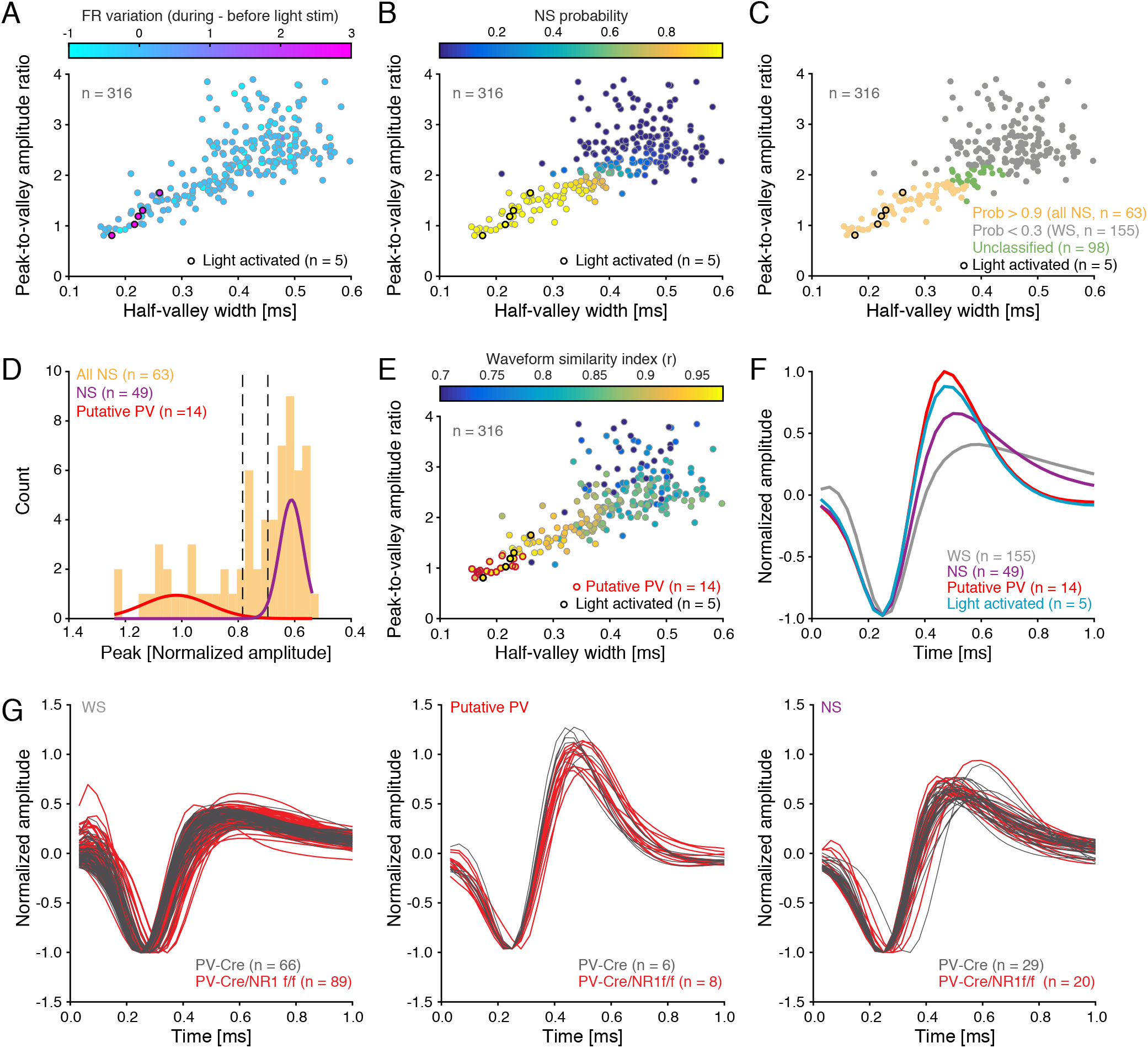
Units classification based on optotagging and spike waveform analysis. **A)** Spike waveform features (mean peak-to-valley amplitude ratio - the ratio between the amplitude of the initial peak and the following trough, and mean half-valley width of spike waveform) for all individual units recorded in mice injected with AAV-DIO-ChR2-mCherry (n = 316 units, PV-Cre (n = 4 mice), PV-Cre/NR1f/f mice (n = 3 mice)). The firing-rate (FR) variation (represented by colorbar) elicited by light application was used to identify light-activated units (n = 5 units). **B)** For the objective classification of units into WS vs NS, a Gaussian mixture model (GMM) was fit to the (1) peak-to-valley amplitude ratio and to the (2) half-valley width of the mean spike waveform of the individual units. This identified the NS probability (represented by colorbar) for the individual units. **C)** Classification of WS and NS units after GMM clustering. Units with a NS probability >0.9 were classified as NS (“all NS”, n = 63), and units with a NS probability <0.3 were classified as WS (n = 155). Units with an intermediate NS probability were unclassified (n = 98 units) and not included in further analysis. **D)** A Gaussian mixture model (GMM) was fit to the peak (normalized amplitude) of the waveform to identify putative PV interneurons within the NS population. NS units with a mean peak amplitude (normalised) >0.8 were classified as putative PV interneurons (n = 14), and units with mean peak amplitude (normalised) <0.3 were classified as NS interneurons (n = 49). **E)** Confirmation of the classification of putative PV interneurons by calculation of the spike waveform similarity index (mean correlation between the unit spike waveform and the average spike waveform of the light-activated PV interneurons) for each unit. The putative PV interneurons identified in (**D**) (n = 14; red outline) exhibit very high spike waveform similarity (*r* > 0.95) with light activated PV interneurons (n = 5; black outline). **F)** Mean normalized spike waveforms for the classified cell types. WS neurons (n = 155 units), putative PV interneurons (n = 14 units), NS interneurons (n = 49 units), and light-activated PV interneurons (n = 5 units). **G)** Mean normalized spike waveforms for the classified WS (left), putative PV (middles), and NS (right) units in PV-Cre (gray) and PV-Cre/NR1f/f (red) mice.

